# Nuclear Import of the HIV-1 Core Precedes Reverse Transcription and Uncoating

**DOI:** 10.1101/2020.03.31.018747

**Authors:** Anastasia Selyutina, Mirjana Persaud, Kyeongeun Lee, Vineet KewalRamani, Felipe Diaz-Griffero

## Abstract

HIV-1 particles contain a core formed by ~1500 capsid protein monomers housing viral RNA. HIV-1 core uncoating---disassembly---is required for infection. HIV-1 reverse transcription (RT) occurs before or during uncoating, but the cellular compartment where RT and uncoating occurs is unknown. Using imaging and biochemical assays to track HIV-1 capsids in nuclei during infection, we demonstrated that higher-order capsid complexes or complete cores containing viral genome are imported into nuclear compartments. Additionally, inhibition of RT that stabilizes the core during infection does not prevent capsid nuclear import; thus, RT may occur in nuclear compartments. We separated infected cells into cytosolic and nuclear fractions to measure RT during infection. Most observable RT intermediates were enriched in nuclear fractions, suggesting that most HIV-1 RT occurs in the nuclear compartment alongside uncoating. Thus, nuclear import precedes RT and uncoating, fundamentally changing our understanding of HIV-1 infection.

## INTRODUCTION

The influence of the physiological state of cells on retroviral replication was initially studied by Rubin and Temin, where they demonstrated that stopping cell division by X-rays or UV light prevented Rous Sarcoma virus replication (Rubin and Temin, 1959; Temin and Rubin, 1959). Subsequent research established the relationship between cell cycle stage and retroviral infection, revealing that diverse retroviruses have different requirements for productive infection (Katz et al., 2005; Suzuki and Craigie, 2007; Yamashita and Emerman, 2004, 2005, 2006; Yamashita et al., 2007). For example, gamma-retroviruses such as murine leukemia virus (MLV) require the host cell to pass through mitosis for productive infection (Lewis and Emerman, 1994; Roe et al., 1993). By contrast, lentiviruses such as HIV-1 show no difference in productive infection when comparing dividing versus non-dividing cells (Lewis et al., 1992). This evidence suggests that lentiviruses have developed specific mechanisms to reach the nuclear compartment.

Although the ability of HIV-1 to enter the nucleus has been attributed to a variety of viral proteins, it is currently believed that the capsid plays a dominant role in the ability of HIV-1 to infect non-dividing cells (Yamashita and Emerman, 2004; Yamashita et al., 2007). HIV-1 particles contain a core formed by approximately 1500 monomers of capsid protein housing the viral RNA (Briggs et al., 2006; Briggs et al., 2004; Briggs et al., 2003). Only approximately 40% of the total capsid protein in the viral particle forms the core (Briggs et al., 2006; Briggs et al., 2004; Briggs et al., 2003). Recent evidence has suggested that the capsid is required for reverse transcription and nuclear import (Hulme et al., 2011; Roa et al., 2012; Yamashita and Emerman, 2004; Yamashita et al., 2007; Yang et al., 2013).

Studies of the rhesus restriction factor TRIM5α led to a major understanding of HIV-1 reverse transcription and uncoating, showing that reverse transcription occurs inside the viral core before or during uncoating, but not after (Diaz-Griffero et al., 2007; Roa et al., 2012; Stremlau et al., 2006). In addition, inhibiting reverse transcription delays HIV-1 uncoating, suggesting that reverse transcription initiation is required for uncoating (Hulme et al., 2011; Yang et al., 2013). These experiments implied that complete or partial integrity of the HIV-1 core is required for reverse transcription and prompted questioning into the true cellular localization of uncoating and reverse transcription. Because integration of HIV-1 viral DNA occurs in the nucleus, it is reasonable to propose that reverse transcription is occurring and/or finishing in the nuclear compartment just prior to integration; however, the precise location where reverse transcription and uncoating occurs is unknown. Nuclear reverse transcription would also require the import of completely or partially assembled capsid into the nucleus, which is currently not understood.

In agreement with a role for the capsid in nuclear import, the HIV-1 core can functionally bind to nucleoporins Nup153 and Nup358 (a.k.a., RANBP2) (Buffone et al., 2018; Di Nunzio et al., 2013; Matreyek et al., 2013). HIV-1 infection of human cells after Nup153 or Nup358 depletion prevents the formation of 2-LTR circles, which indirectly indicates that the pre-integration complex has not reached the nucleus (Butler et al., 2001). Additionally, the use of imaging has revealed the presence of capsid in the nucleus (Bejarano et al., 2019; Chin et al., 2015; Hulme et al., 2015; Peng et al., 2014). Although tracking single particles by fluorescence microscopy has detected HIV-1 replication complexes in the nucleus (Bejarano et al., 2019; Burdick et al., 2017; Francis and Melikyan, 2018), the total amount of capsid in the nucleus and its importance for infection have not been studied. This work therefore sought to explore the importance of nuclear capsid for productive infection.

Using biochemical assays to track the fraction of capsid in the nucleus during infection, we demonstrate that at any given time during the early phase of HIV-1 infection, approximately 20-30% of total capsid is in the nucleus. Remarkably, we found that assembled capsid complexes are imported into the nuclear compartment, suggesting that parts of the core or complete cores containing the viral genome are transported into the nucleus. In agreement with these findings, we found that inhibiting reverse transcription, which stabilizes the core during infection, does not prevent capsid import into the nucleus, indicating that reverse transcription is occurring in the nuclear compartment. To test this hypothesis, we separated infected cells into cytosolic and nuclear fractions to measure the steps of reverse transcription over time. Our results reveal that most of the observable reverse transcription intermediates are enriched in the nuclear compartment, inferring that most HIV-1 reverse transcription occurs in the nuclear compartment. Because reverse transcription occurs in the nucleus, these experiments also imply that uncoating proceeds in the nuclear compartment. These findings fundamentally change our understanding of HIV-1 infection and conclude that nuclear import precedes reverse transcription and uncoating.

## RESULTS

### Biochemical assay to detect capsid in the nuclei of HIV-1 infected human cells

Tracking single HIV-1 particles by microscopy permitted the detection of capsids in nuclei (Bejarano et al., 2019; Chin et al., 2015; Hulme et al., 2015; Peng et al., 2014); however, the contribution of HIV-1 capsid to nuclear import, reverse transcription, and productive infection is not clear. To study the role of capsid in the nucleus, we developed a methodology to isolate the nuclear content of HIV-1-infected cells to measure the amount of capsid (Figure 1A). This methodology consists of performing synchronized infections of human cells with HIV-1 using an MOI = 2 and fractionating cells at different time points after infection (see Methods). Separation of nuclear and cytoplasm fractions are carried out through the use of buffers containing detergent and different salt concentrations. Nuclear and cytosolic fractions are identified by western blotting using anti-Nopp140 and anti-GAPDH antibodies, respectively, then analyzed for HIV-1 capsid by western blotting using anti-p24 antibodies (Figure 1A).

**Figure 1.**
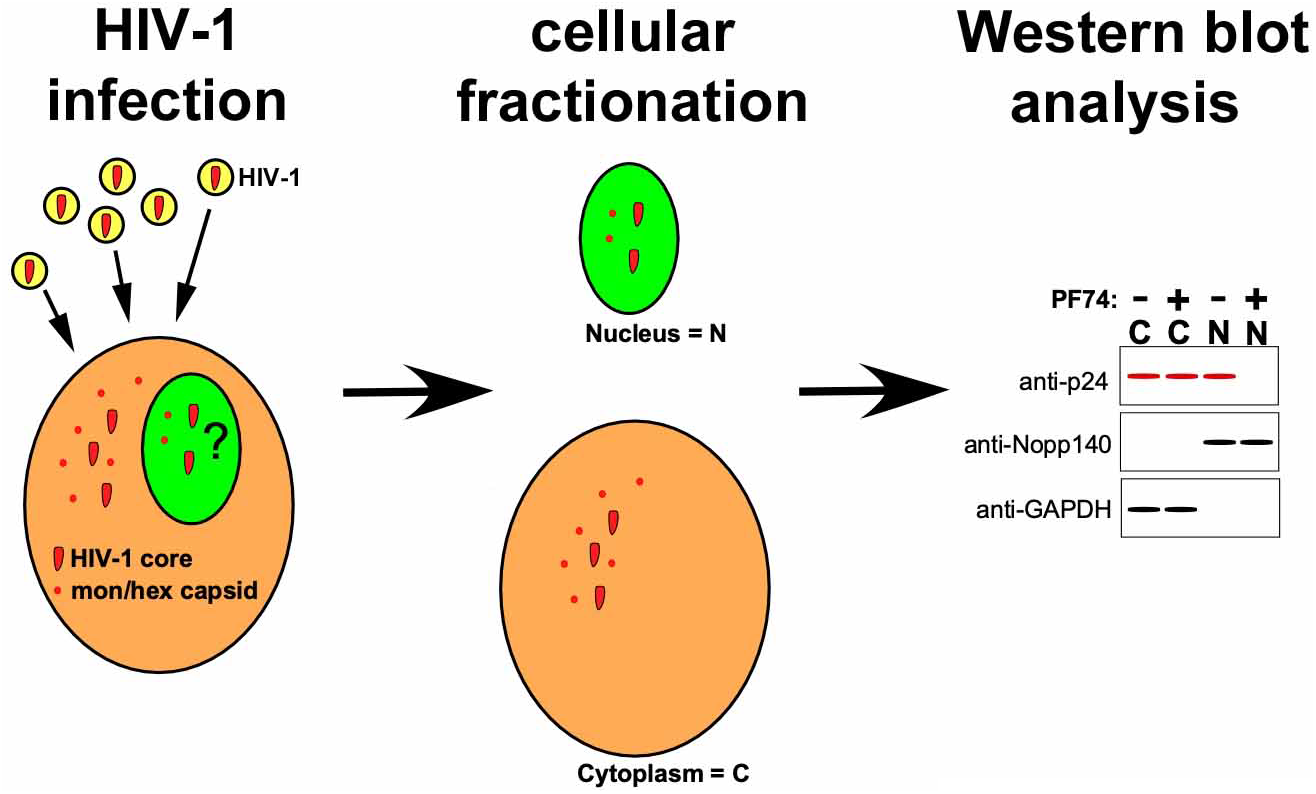
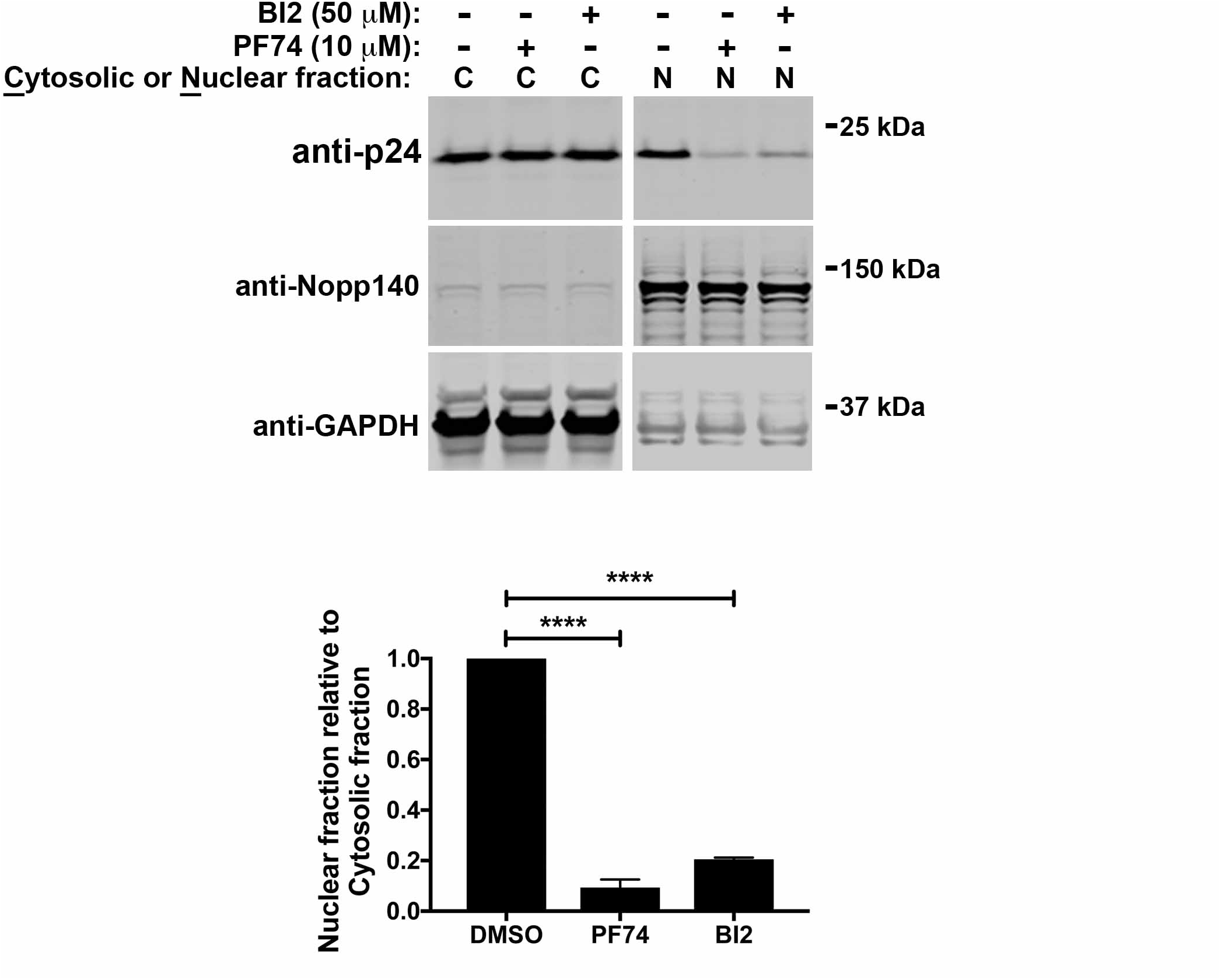
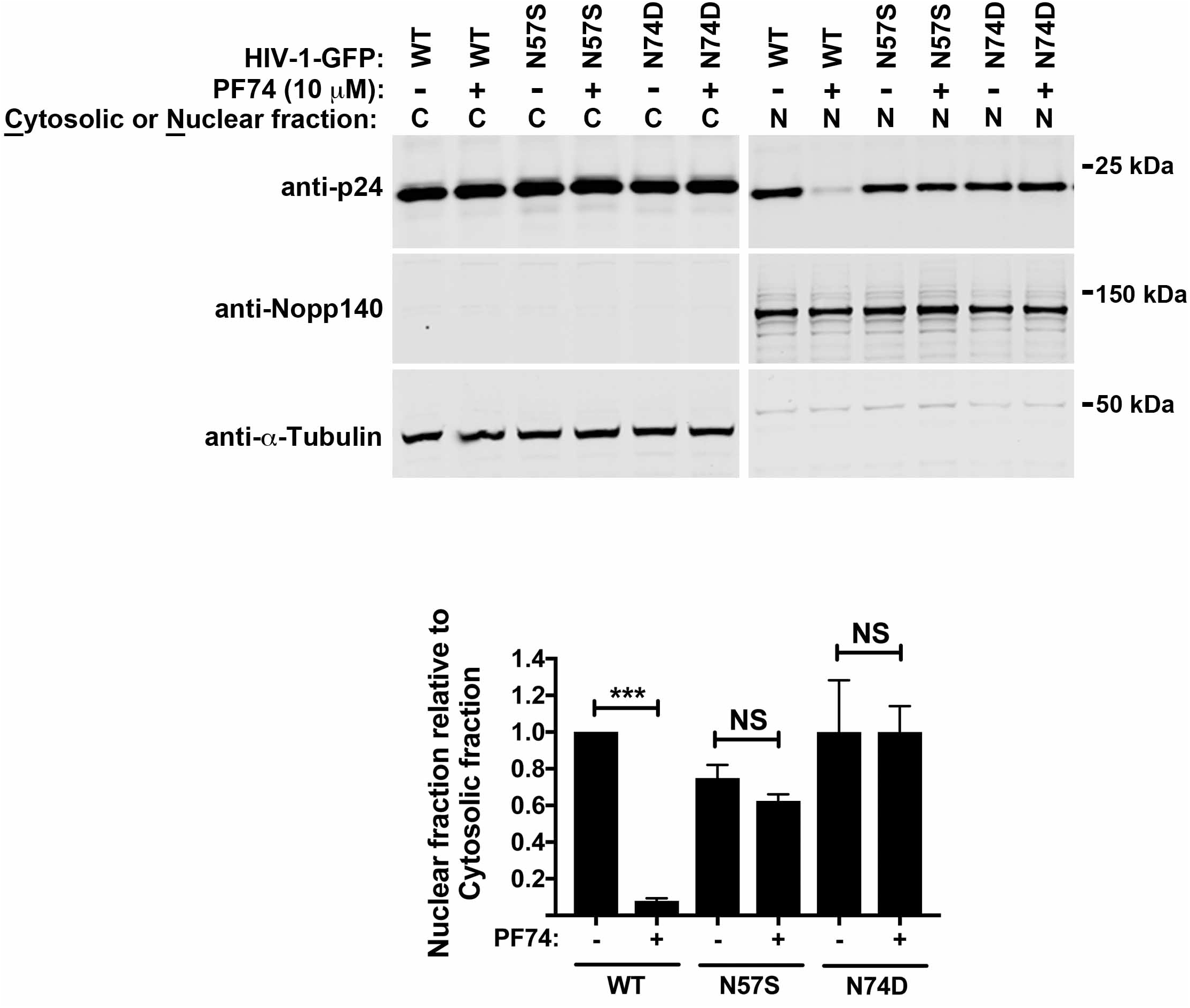
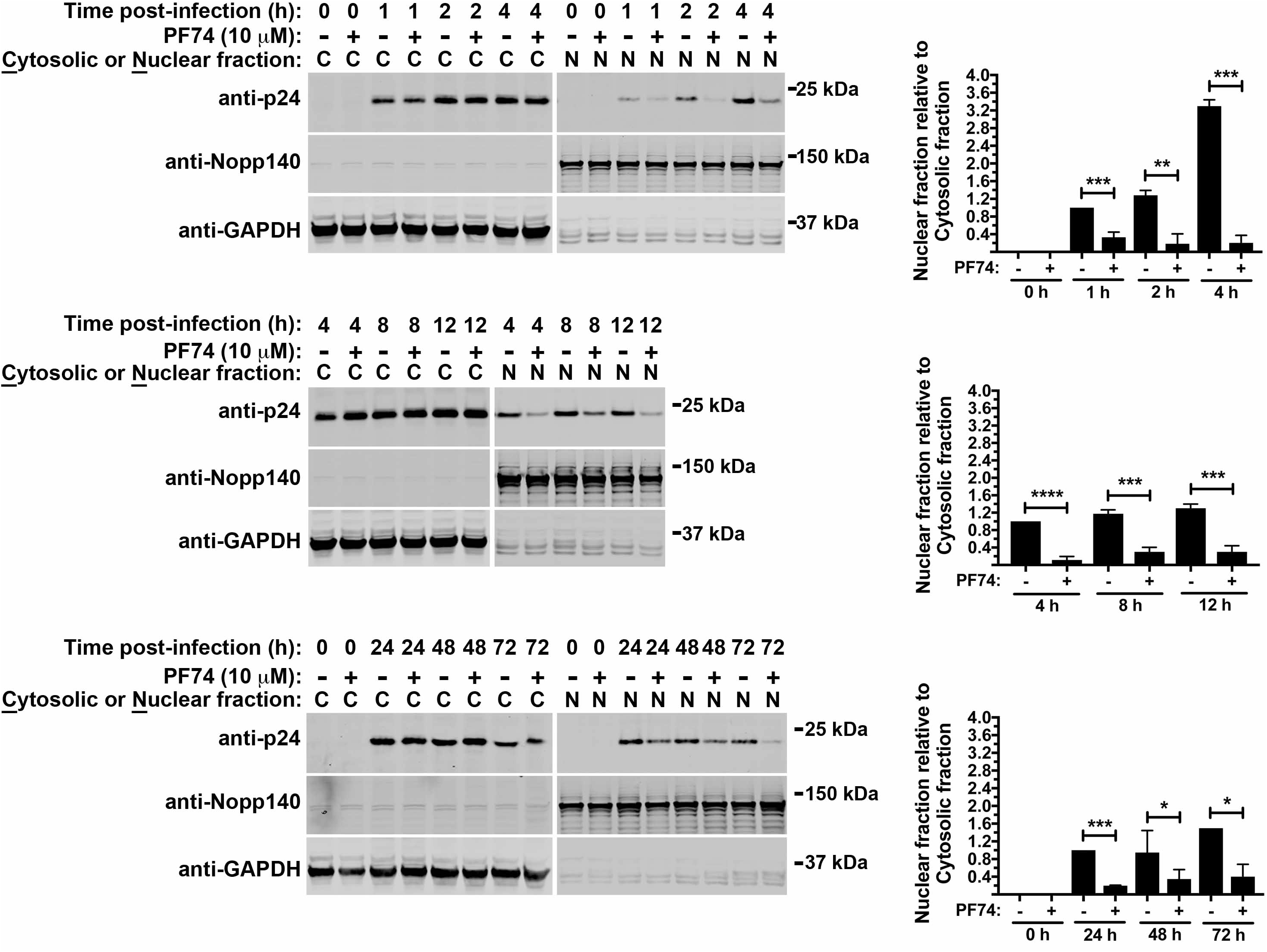
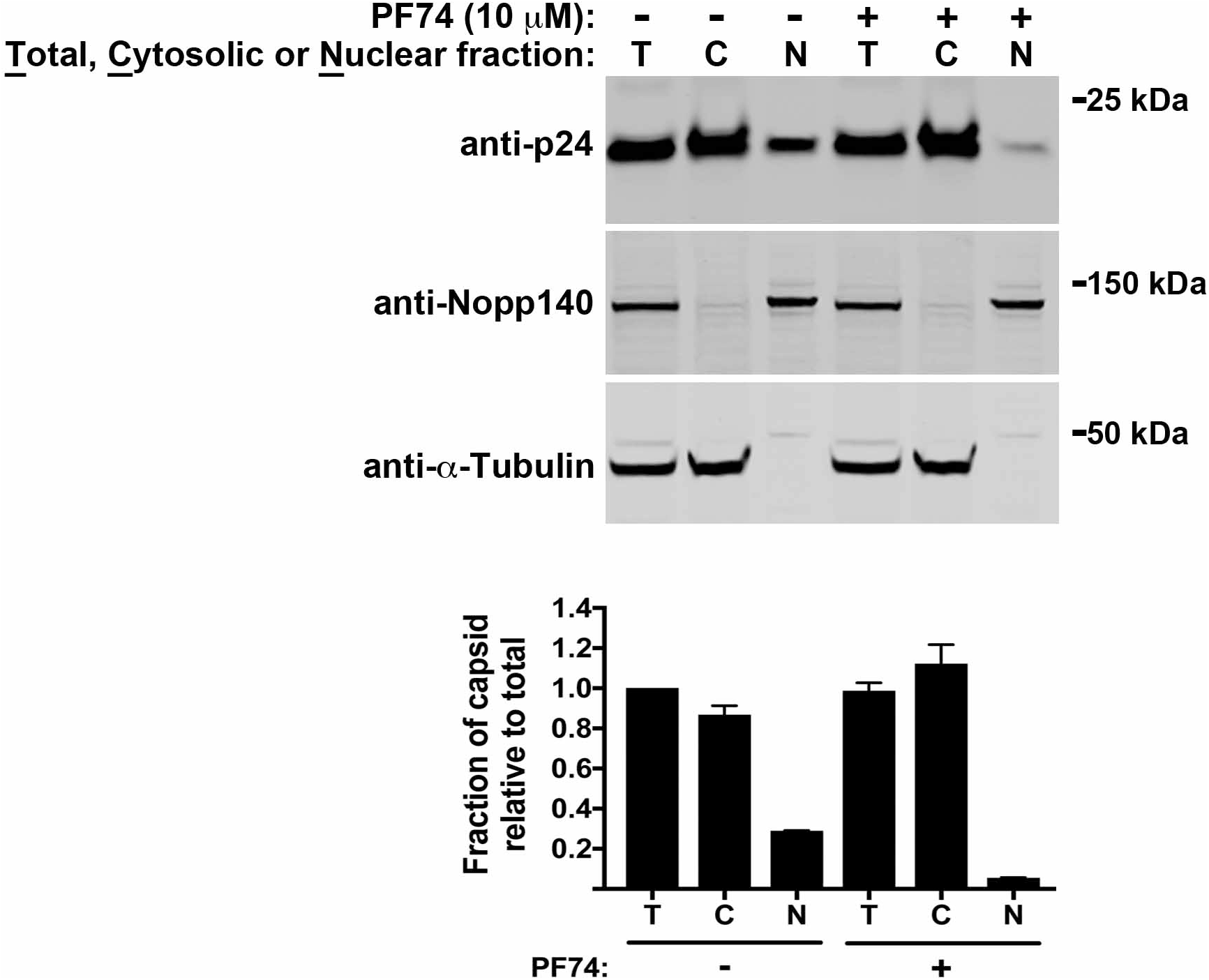
Biochemical assay to detect capsid in the nucleus during HIV-1 infection. **(A)** Schematic representation of subcellular fractionation. Human cells infected with HIV-1 in the presence or absence of PF74 are separated into cytosolic and nuclear fractions using detergents and buffers with different salt concentrations. The small HIV-1 inhibitor PF74 is used as a control since it prevents the nuclear import of capsid. Subsequently, fractions are analyzed by western blotting using anti-p24 antibodies. To verify the origin and purity of cellular fractions, western blotting was performed using anti-Nopp140 and anti-GAPDH antibodies, which are nuclear and cytosolic markers, respectively. **(B)** Treatment with PF74 or BI-2 prevents capsid import into the nucleus. A549 cells were infected with HIV-1-GFP at MOI = 2 for 8 h in the presence of 10 μM PF74, 50 μM BI2, or DMSO as a vehicle control. Subsequently, cells were separated into nuclear and cytosolic fractions and analyzed for capsid content by western blotting using anti-p24 antibodies. Nopp140 and GAPDH detection by western blotting was used as a marker for nuclear and cytosolic content, respectively. **(C)** PF74 does not affect capsid nuclear import by mutant HIV-1-N57S-GFP and HIV-1-N74D-GFP viruses. A549 cells were infected with p24-normalized HIV-1-GFP, HIV-1-N57S-GFP, or HIV-1-N74D-GFP viruses in the presence of 10 μM PF74 or DMSO as a vehicle control for 8 h. Cells were then separated into nuclear and cytosolic fractions and analyzed for capsid content by western blotting using anti-p24 antibodies. Detection of Nopp140 and tubulin by western blotting was used as a marker for nuclear and cytosolic content, respectively. **(D)** Nuclear import of capsid during infection. A549 cells were infected with HIV-1-GFP at MOI = 2 in the presence of 10 μM PF74 or DMSO as a vehicle control. At the indicated times, cells were separated into nuclear and cytosolic fractions and analyzed for capsid content by western blotting using anti-p24 antibodies. Detection of Nopp140 and GAPDH by western blotting was used as a marker for nuclear and cytosolic content, respectively. Experiments were repeated at least three times and a representative figure is shown. The ratio of Nuclear to Cytosolic capsid for three independent experiments with standard deviations is shown. **(E)** Relative amounts of capsid protein in total (T), cytosolic (C), and nuclear (N) fractions. A549 cells were infected with HIV-1-GFP at a MOI = 2 in the presence of 10 μM PF74 or DMSO as a vehicle control for 8 h. Cells were separated into nuclear and cytosolic fractions. Total, cytosolic, and nuclear fractions were analyzed by western blotting using anti-p24, anti-Nopp140, and anti-GAPDH antibodies. Experiments were repeated at least three times and a representative figure is shown. The amount of capsid relative to the total for three independent experiments with standard deviations is shown. * indicates P-value < 0.005, ** indicates P-value < 0.001, *** indicates P-value < 0.0005, **** indicates P-value < 0.0001, NS indicates not significant as determined by using the unpaired t-test.

Because the use of the small molecules PF74 and BI-2 blocks HIV-1 infection before the production of 2-LTR circles and prevents Nup153 binding to capsid (Balasubramaniam et al., 2019; Buffone et al., 2018; Fricke et al., 2014a), we decided to test whether these compounds had an effect on the import of capsid into the nucleus during infection. To this end, we performed synchronized infections of human A549 cells using HIV-1-GFP viruses (MOI = 2) for 8 h. 10 μM PF74 or 50 μM BI-2 prevented the import of HIV-1 capsid into the nucleus (Figure 1B). These results agree with previous findings demonstrating that PF74 and BI-2 prevent 2-LTR circle formation (Fricke et al., 2014a), which is interpreted as an inhibition of nuclear import of the HIV-1 replication complex (Butler et al., 2001). PF74 inhibition of HIV-1 capsid import into the nucleus was utilized as a control for this assay.

To test the specificity of this assay for measuring the nuclear import of capsid, we measured capsid nuclear import of HIV-1 virus bearing the capsid changes N57S and N74D that result in viral resistance to PF74 (Buffone et al., 2018; Saito et al., 2016). To this end, we fractionated human A549 cells infected with HIV-1-N57S and HIV-1-N74D viruses in the presence of PF74 for 8 h. PF74 did not affect the nuclear import of capsid from HIV-1-N57S and HIV-1-N74D viruses (Figure 1C), which are resistant to PF74 (Buffone et al., 2018; Saito et al., 2016). Overall these results demonstrated that our assay specifically measures the import of HIV-1 capsid into the nucleus and that infection correlates with capsid nuclear import.

We used this assay to investigate the earliest time necessary for capsid detection in the nucleus by fractionating A549 cells infected for the indicated times using wild-type HIV-1. As shown Figure 1D, we detected capsid in the nucleus as early as 1 h postinfection, which appears to increase over time and peak at 8 h post-infection. As expected, the amount of nuclear capsid 1 h post-infection was sensitive to PF74. The peak of capsid in the nucleus seems to coincide with the peak of reverse transcription as measured by real-time PCR, which is 8-10 h post-infection. After 12 h post-infection, the amount of nuclear capsid starts decreasing. Collectively, these experiments show that the amount of nuclear capsid increases as infection proceeds over time.

Our novel assay biochemically demonstrated the presence of capsid in the nucleus, raising questions about the amount of total capsid present in the nucleus. Using this semi-quantitative assay, we challenged human A549 cells with wild-type HIV-1 virus for 8 h. Capsid levels were determined by fractionating cells into total (T), nuclear (N), and cytoplasmic (C) fractions. Approximately 20-30% of total capsid corresponded with the nuclear fraction (Figure 1E). These results suggest that the fraction of capsid imported into the nucleus corresponds with approximately the amount of capsid that is forming HIV-1 cores in the viral particle (Briggs et al., 2006; Briggs et al., 2004; Briggs et al., 2003). Although human A549 cells are a reliable model to study HIV-1 infection, we next tested our nuclear fractionation assay using the T-cell line MOLT3. As shown in Figure S1, PF74 prevented the import of HIV-1 capsid into the nucleus of T cells.

To corroborate these biochemical findings, we performed similar infections and imaged the viral capsid using confocal microscopy in the presence of PF74. For the purpose of imaging, the infections were synchronized. As shown in Figure S2, PF74 prevented the entry of capsid into the nucleus. These results agree with our biochemical observations showing that PF74 prevents the import of HIV-1 capsid into the nucleus.

### Aggregation of CPSF6 in the nucleus during HIV-1 infection requires intact nuclear capsid

Infection of human cells by HIV-1 triggers a change in the nuclear staining pattern of CPSF6 from diffuse to speckled at 8 or 16 h post-infection when compared to mock controls (Figure 2). To functionally establish the presence of capsid in the nucleus, we investigated whether nuclear import of HIV-1 capsid is required for CPSF6 aggregation into a speckled distribution, human HeLa cells were challenged with wildtype and mutant HIV-1 viruses in the presence of PF74 using an MOI = ~2 for 8 or 16 h. Subsequently, cells were fixed and stained for CPSF6 and capsid, and cellular nuclei were counterstained with DAPI. As shown in Figure 2, HIV-1 infection induced CPSF6 speckling in the nuclear compartment at 8- and 16-h post-infection. By contrast, uninfected cells showed a diffuse staining pattern for CPSF6. Inhibition of capsid nuclear import with PF74 during HIV-1 infection resulted in a diffuse staining of CPSF6, suggesting that the presence of capsid in the nucleus is required for CPSF6 to converge into nuclear speckles. We next investigated whether nuclear capsid is sufficient for CPSF6 speckle formation by challenging HeLa cells using HIV-1-N74D and HIV-1-N57S viruses, which are viruses that import capsid into the nucleus during infection without capsid interaction with CPSF6 (Buffone et al., 2018; Lee et al., 2010). Although HIV-1-N74D and HIV-1-N57S viruses underwent normal capsid import (see Figure 1C), CPSF6 showed a diffuse staining pattern (Figure 2). These experiments suggest that capsid interaction with CPSF6 is required for the formation of CPSF6 nuclear speckles. Overall, these experiments show that the formation of nuclear CPSF6 speckles requires the interaction of CPSF6 with capsid in the nucleus.

**Figure 2.**
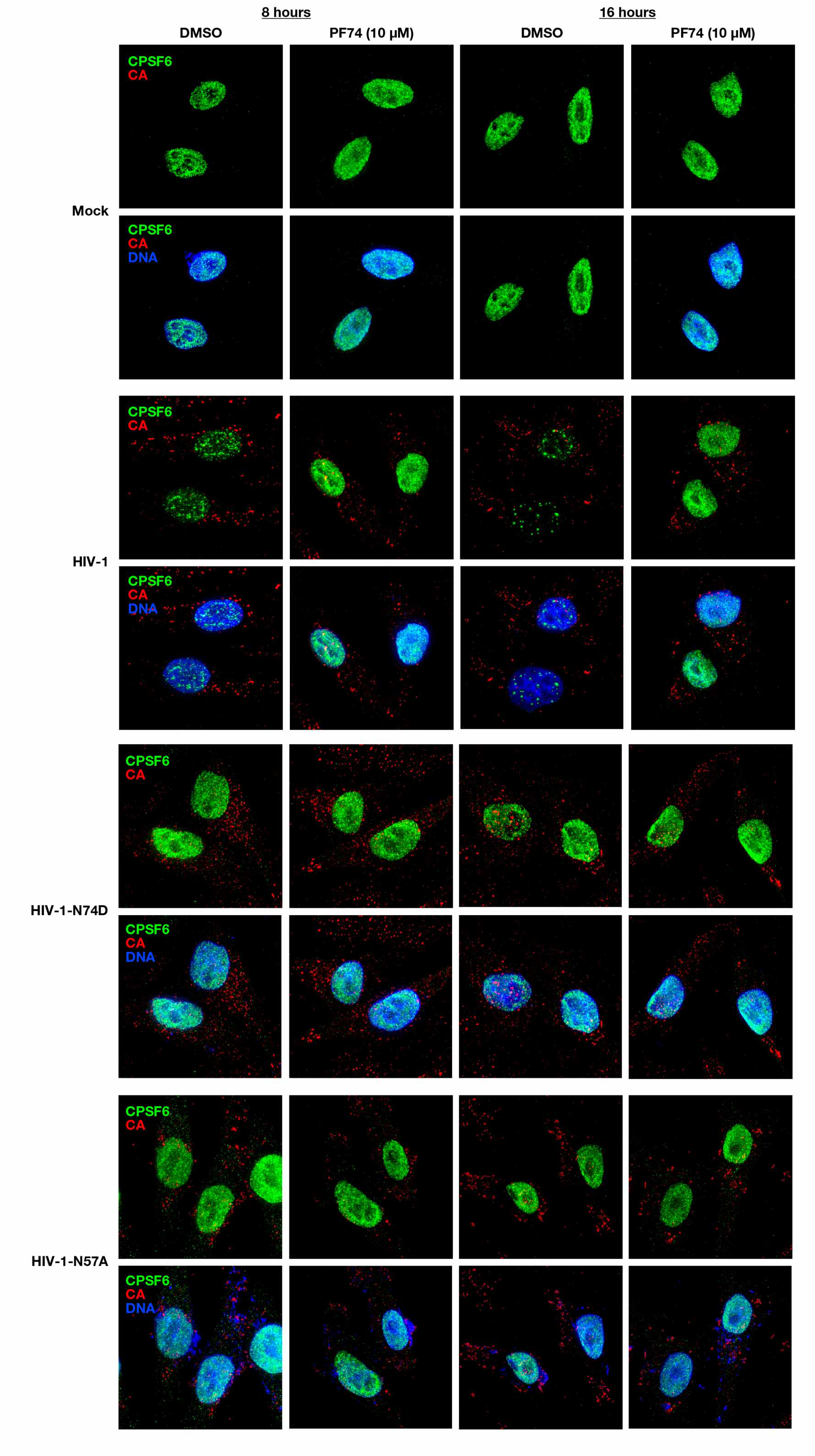
CPSF6 aggregation in the nucleus during HIV-1 infection requires interaction with nuclear capsid. HeLa cells were infected with wild-type HIV-1, mutant HIV-1-N74D, mutant HIV-1-N57A viruses at MOI = X in the presence of 10 μM PF74 or DMSO as a vehicle control. After incubation for 8 or 16 h, cells were fixed and stained for CPSF6 and HIV-1 capsid (CA). Cellular nuclei were counterstained with DAPI. Experiments were repeated at least three times and a representative figure is shown.

### Assembled capsid complexes are imported into the nuclear compartment during HIV-1 infection

Capsid import into the nucleus during infection may be occurring by different mechanisms: (1) monomeric capsid is imported into the nucleus, (2) assembled capsid complexes are imported into the nucleus, or (3) both. To test whether assembled HIV-1 capsid complexes are imported into the nucleus, we measured nuclear capsid in cells expressing the restriction factors TRIMCyp and rhesus TRIM5α (TRIM5α_rh_), which accelerate HIV-1 uncoating in the cytoplasm of infected cells (Diaz-Griffero et al., 2007; Stremlau et al., 2006). Accelerated uncoating causes the disassembly of higher order capsid complexes, such as the core. To this end, we challenged human A549 cells expressing TRIMCyp or TRIM5α_rh_ with HIV-1 at a MOI = 2 for 8 h. TRIMCyp and TRIM5α_rh_, which potently blocked HIV-1 infection (Figure S3), prevented HIV-1 capsid import into the nucleus (Figure 3A). These experiments demonstrate that the presence of capsid in the nucleus results from the import of larger capsid complexes, as has been shown for other viruses (Cohen et al., 2011; Fay and Pante, 2015; Whittaker et al., 2000). These larger capsid complexes may be complete cores or pieces of the core that are associated with the viral genome.

**Figure 3.**
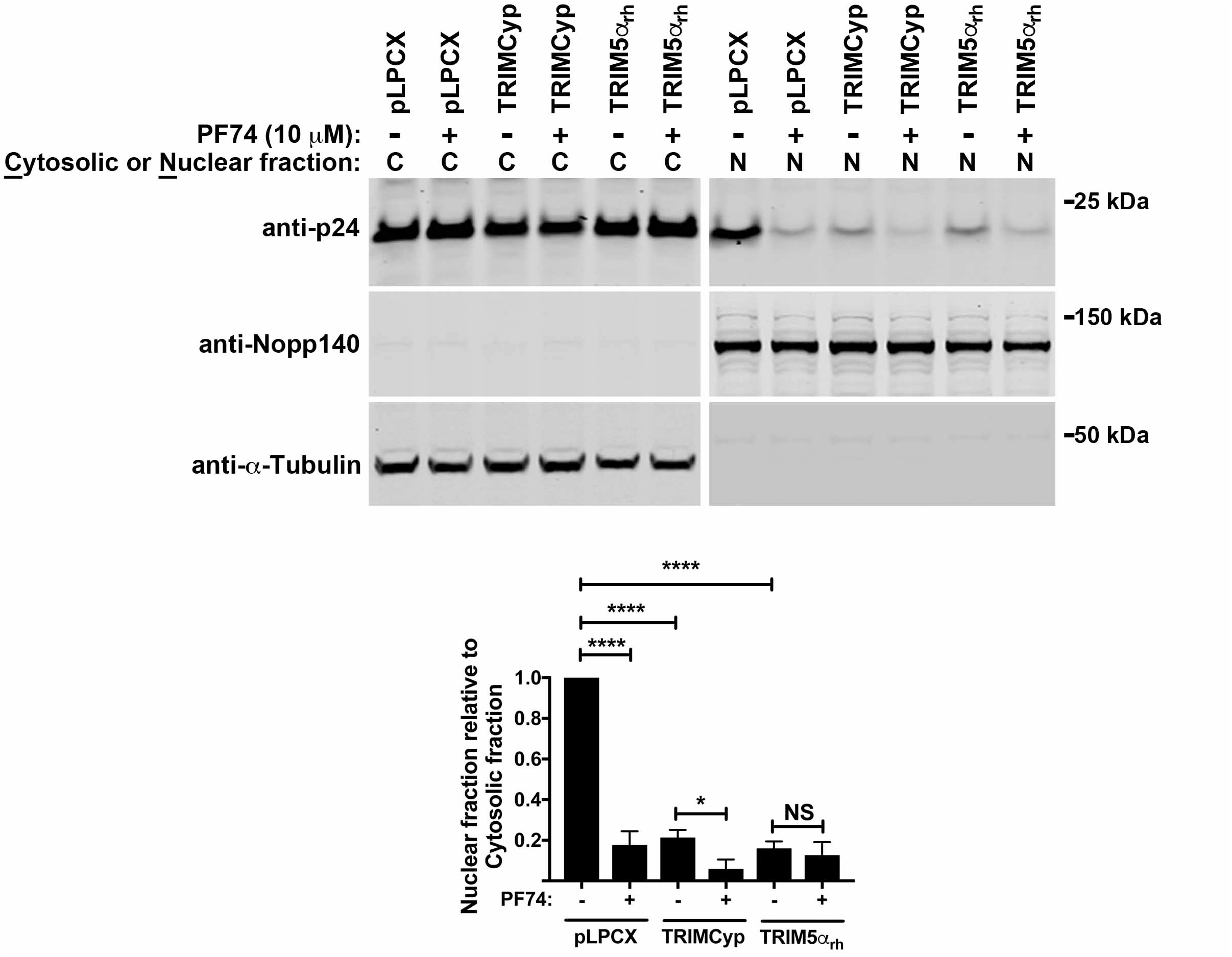
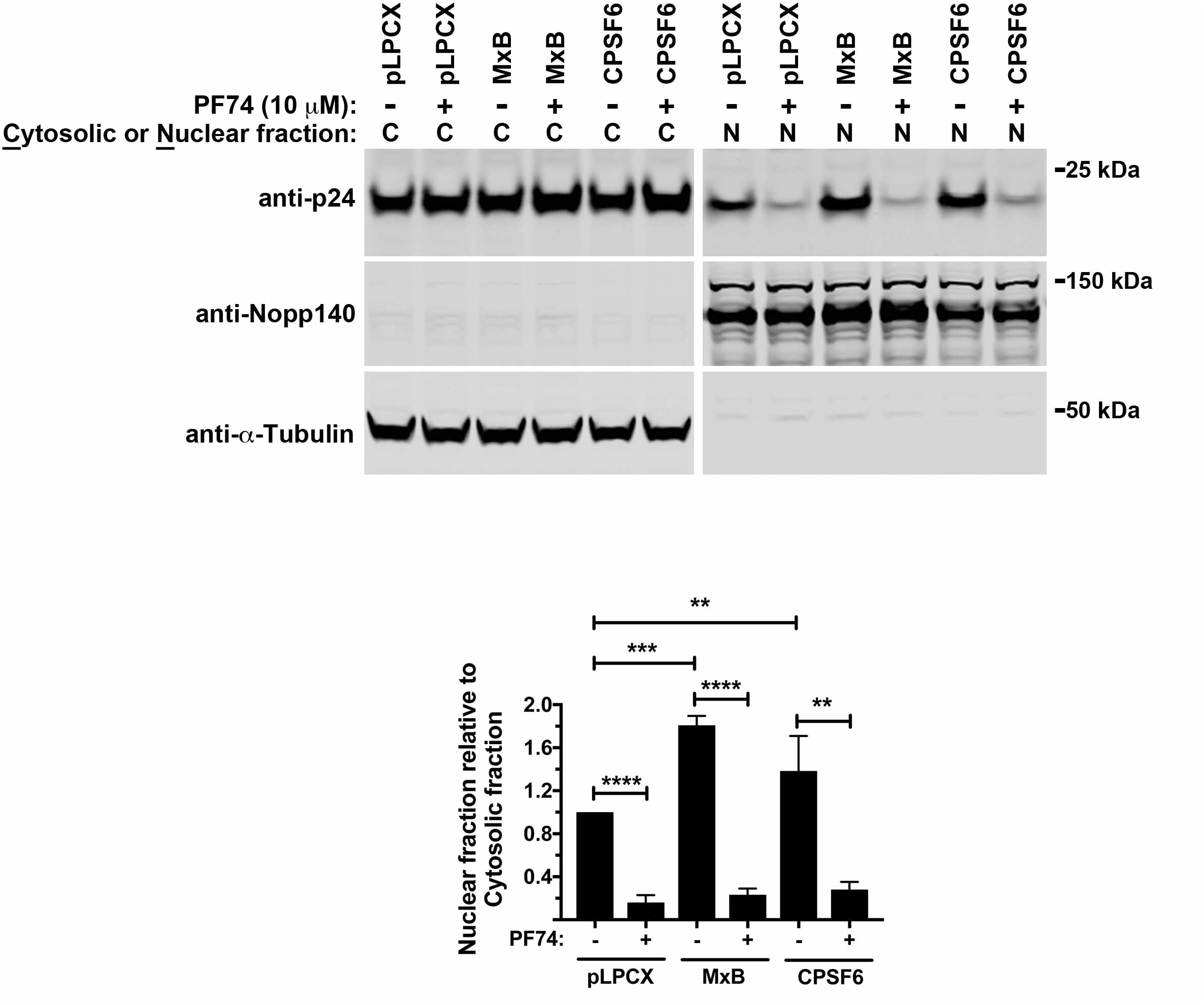
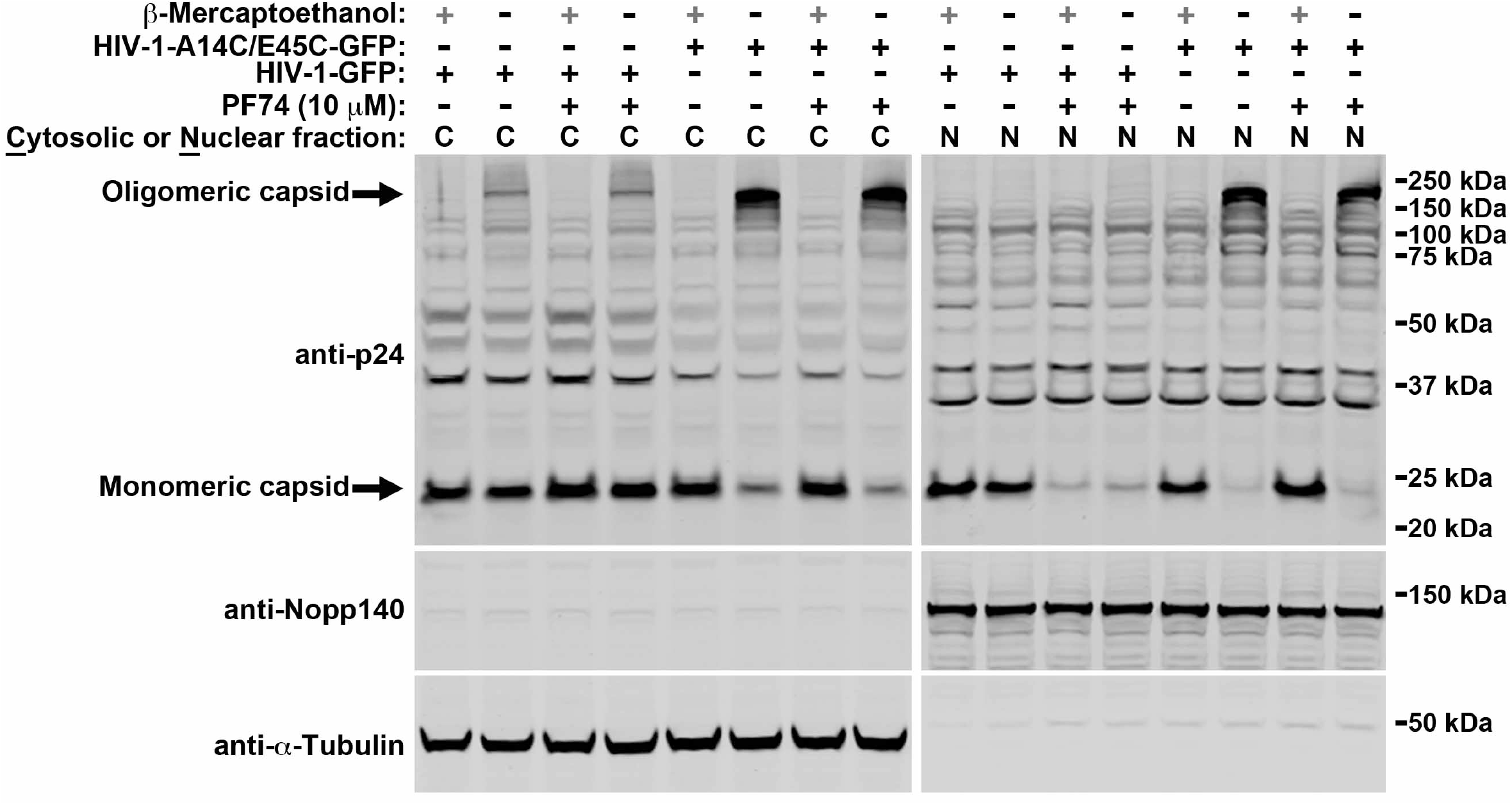
Assembled capsid complexes are imported into the nuclear compartment during HIV-1 infection. Human cells stably expressing rhesus TRIM5α_rh_ **(A)**, owl monkey TRIMCyp **(A)**, human NES-CPSF6(1-358) **(B)**, human MxB proteins **(B)**, or empty pLPCX vector, were infected with wild-type HIV-1-GFP at MOI = 2 in the presence of 10 μM PF74 or DMSO as a vehicle control for 8 h. Cells were separated into nuclear and cytosolic fractions and analyzed for capsid content by western blotting using anti-p24 antibodies. Detection of Nopp140 and tubulin by western blotting was used as a marker for nuclear and cytosolic content, respectively. Experiments were repeated at least three times and a representative figure is shown. The ratio of Nuclear to Cytosolic capsid for three independent experiments with standard deviations is shown. ** indicates P-value < 0.001, *** indicates P-value < 0.0005, **** indicates P-value < 0.0001 as determined by using the unpaired t-test. **(C)** Human A549 cells were infected using p24-normalized amounts of wild-type HIV-1-GFP and mutant HIV-1-A14C/E45C-GFP viruses (virus amount corresponded to wild-type MOI = 2) in the presence of 10 μM PF74 or DMSO as a vehicle control for 8 h. Cells were separated into nuclear and cytosolic fractions and analyzed for capsid content by western blotting using anti-p24 antibodies in the presence or absence of the reducing agent β-mercaptoethanol. Detection of Nopp140 and tubulin by western blotting was used as a marker for nuclear and cytosolic content, respectively. Experiments were repeated three times(Figure S6) and a representative figure is shown.

We explored whether factors that stabilize the HIV-1 core during infection affect the presence of capsid in the nucleus in cells expressing MxB or CPSF6 that contains the nuclear export signal of PKI_α_ (NES-CPSF6) (Fricke et al., 2013). These proteins prevent uncoating by stabilizing the HIV-1 core during infection (De Iaco et al., 2013; Fricke et al., 2013; Fricke et al., 2014b). Expression of MxB or NES-CPSF6, which potently restricts HIV-1 (Figure S3), did not affect the ability of the HIV-1 capsid to reach the nuclear compartment (Figure 3B). These results demonstrate that stabilized HIV-1 capsid complexes, which may be intact cores or slightly uncoated cores, are imported into the nuclear compartment, which agrees with our previous findings suggesting that large capsid complexes or complete cores are the source of nuclear capsid in our assay.

Experiments using our nuclear import functional assay suggested that assembled capsids are transported into the nucleus; however, the western blots cannot distinguish between assembled or disassembled capsid within the nucleus. To directly test whether large assembled capsid complexes can be imported into the nucleus using an HIV-1 core resistant to disassembly, we produced an HIV-1 virus with a stabilized capsid, taking advantage of the capsid mutations A14C/E45C that stabilize purified capsid hexamers *in vitro* through disulfide bridges between monomers of hexamers (Pornillos et al., 2010; Selyutina et al., 2018). This virus lacked defects in viral budding, maturation, and reverse transcription; however, HIV-1-A14C/E45C virus was incapable of forming 2-LTR circles and productive infection (Figure S4). In addition, the core of HIV-1-A14C/E45C virus showed greater stability when compared to wild-type in our capsid assay (Figure S5). Human A549 cells were challenged using HIV-1-A14C/E45C virus, and nuclear fractions were isolated 8 h post-infection. To distinguish assembled from disassembled capsid, the nuclear fractions were analyzed in the presence or absence of the reducing agent β-mercaptoethanol. Remarkably, most of the HIV-1-A14C/E45C capsid in the nuclear fraction was in the assembled form since its migration by SDS-PAGE was sensitive to β-mercaptoethanol (Figure 3C and Figure S6). These results illustrate that large, assembled capsid complexes are imported into the nuclear compartment. Additionally, nuclear import of HIV-1-A14C/E45C capsid was not sensitive to PF74, which may be due to the enhanced stability of HIV-1-A14C/E45C cores. This corresponds with our earlier findings inferring that the observed nuclear capsids are a result of importing assembled capsid complexes, such as complete cores.

### Reverse transcription inhibition does not affect nuclear capsid levels during HIV-1 infection

Our studies demonstrate that large assembled capsid complexes are imported into the nuclear compartment. Previous evidence has shown that genetically or pharmacologically inhibition of reverse transcription enhances HIV-1 core stability (Hulme et al., 2011; Yang et al., 2013). Furthermore, acceleration of uncoating, or the peeling of monomeric capsids from the core, correlates with the inhibition of reverse transcription, suggesting that reverse transcription occurs before or during uncoating (Arfi et al., 2009; Roa et al., 2011). Given these observations, we decided to test whether inhibiting reverse transcription affects nuclear import by infecting A549 cells with HIV-1 in the presence of the reverse transcription inhibitors nevirapine and AZT. Pharmacological inhibition of reverse transcription did not affect capsid import into the nucleus during infection, nor did genetic ablation of reverse transcription using HIV-1 virus bearing the mutation D185N in the reverse transcriptase enzyme (Yang et al., 2013) (Figure 4A). These results are in agreement with the notion that large assembled capsid complexes are imported into the nucleus while reverse transcription is occurring.

**Figure 4.**
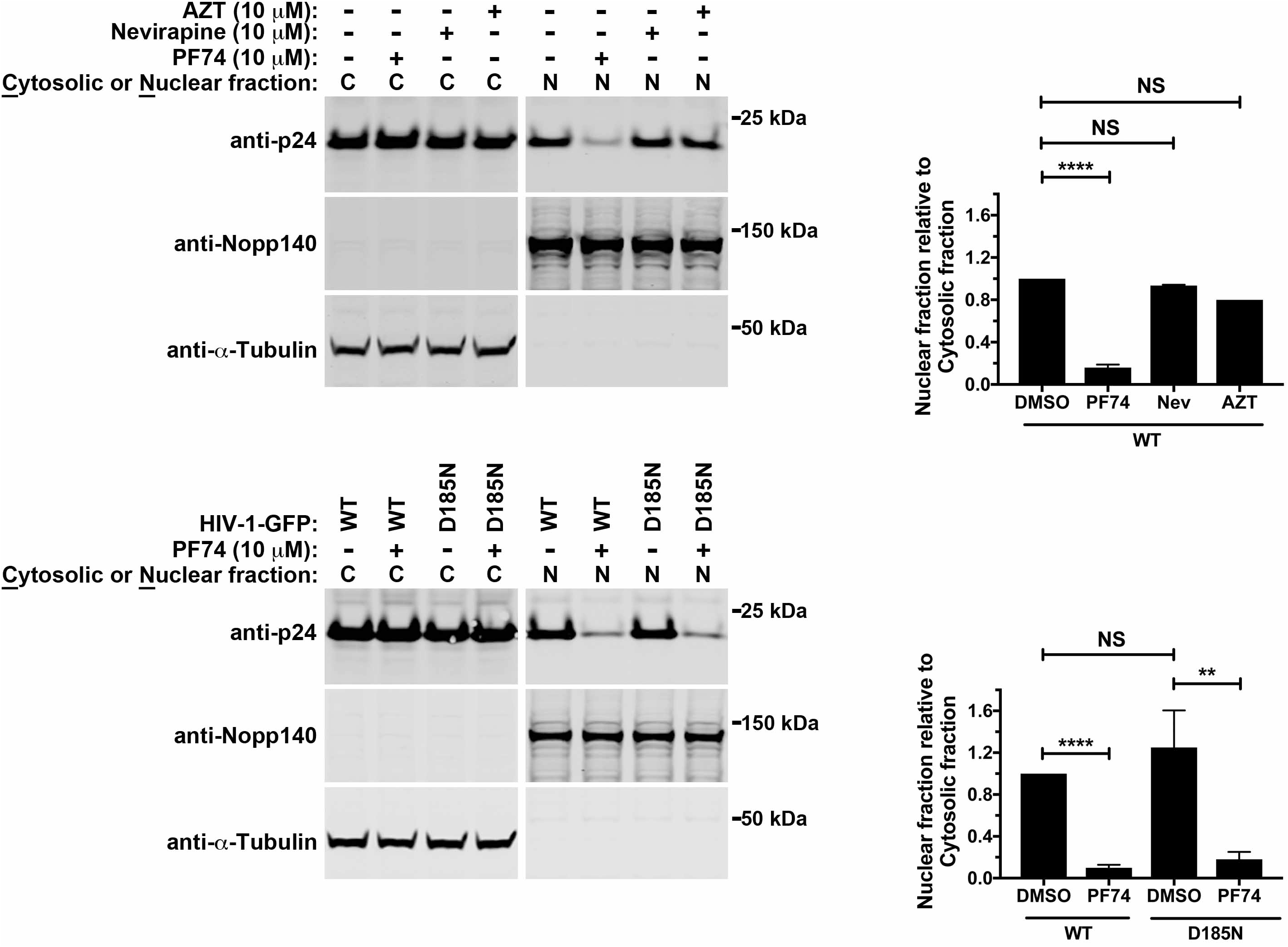
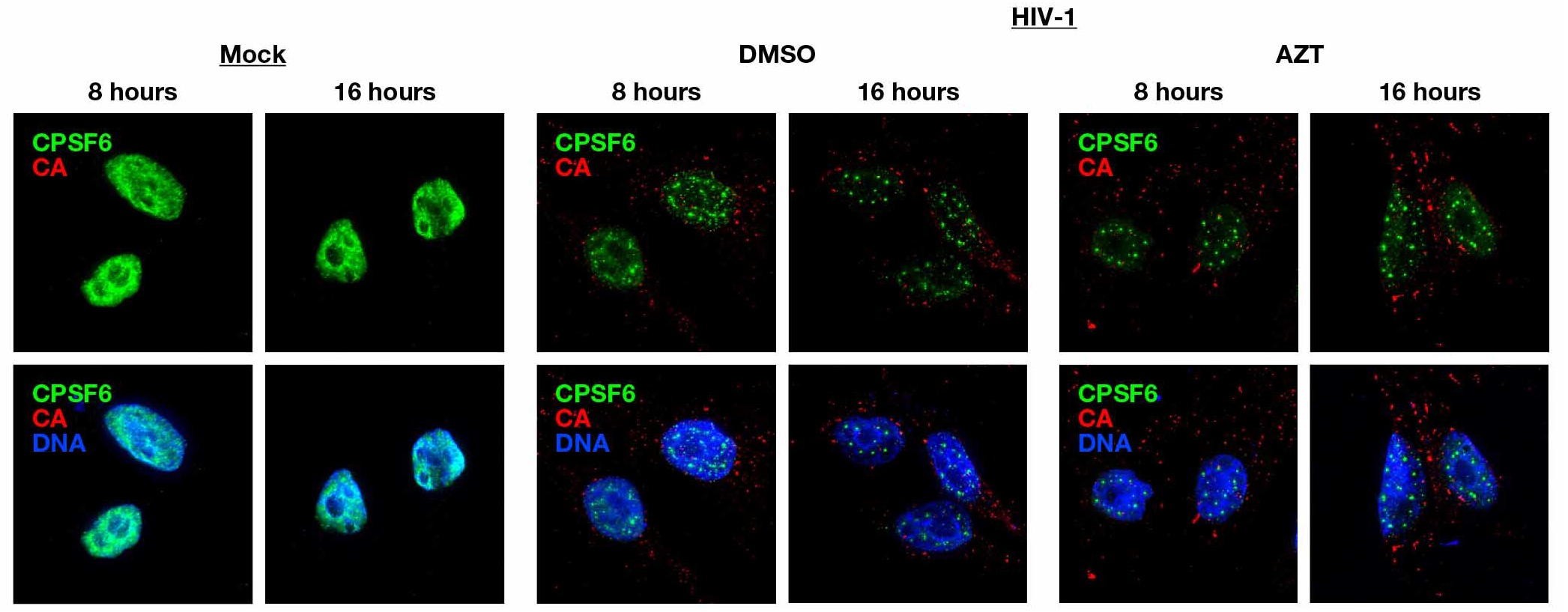
Inhibiting reverse transcription does not affect the levels of nuclear capsid during HIV-1 infection. **(A)** Human A549 cells were infected with wild-type HIV-1-GFP at MOI = 2 in the presence of 10 μM PF74, 10 μM AZT, 10 μM nevirapine, or DMSO as a vehicle control for 8 h. Cells were separated into nuclear and cytosolic fractions and analyzed for capsid content by western blotting using anti-p24 antibodies. Detection of Nopp140 and tubulin by western blotting was used as a marker for nuclear and cytosolic content, respectively. Similar fractionation experiments were performed in cells infected with p24-normalized HIV-1-GFP and mutant HIV-1-D185N viruses. The D185N mutation results in a modified reverse transcriptase lacking enzymatic activity. Experiments were repeated at least three times and a representative figure is shown. The ratio of Nuclear to Cytosolic capsid for three independent experiments with standard deviations is shown. ** indicates P-value < 0.001, **** indicates P-value < 0.0001, NS indicates not significant as determined by using the unpaired t-test. **(B)** HeLa cells were infected with wild-type HIV-1 at MOI = ~2 in the presence of 10 μM AZT or DMSO as a vehicle control. After incubation for 8 or 16 h, cells were fixed and stained for CPSF6 and HIV-1 capsid (CA). Cellular nuclei were counterstained with DAPI. Experiments were repeated at least three times and a representative figure is shown.

To evaluate the functionality of the nuclear HIV-1 capsid during reverse transcription inhibition, we monitored the formation of CPSF6 speckling by challenging HeLa cells with HIV-1 in the presence of AZT for 8 and 16 h. Cells were fixed and stained for CPSF6 and capsid, and nuclei were counterstained using DAPI. As shown in Figure 4B, infection in the presence or absence of AZT resulted in the formation of CPSF6 speckles, showing that nuclear capsid is functional in the presence of reverse transcription inhibitors since it behaves similarly to the non-inhibited virus.

### HIV-1 reverse transcription occurs in the nuclear compartment

Our results have indicated that large assembled capsid complexes can enter the nuclear compartment even during reverse transcription inhibition, supporting the hypothesis that reverse transcription occurs in the nuclear compartment. To further validate these observations, Cf2Th cells were challenged with HIV-1 virus for 2, 4, and 8 h, after which nuclear and cytosolic fractions were concurrently analyzed for capsid protein content (Figure 5A and Figure S7) and monitored using real-time PCR for early products, minus strand transfer, intermediate products, and late reverse transcribed products from equal amounts of fractions (Figure 5B and Figure S7). As controls, the reverse transcription inhibitor nevirapine and mock-infected cells were utilized. All reverse transcription intermediates tested showed higher capsid presence in the nuclear fraction when compared to the cytosolic fraction. These results indicate that HIV-1 reverse transcription occurs in the nuclear compartment.

**Figure 5.**
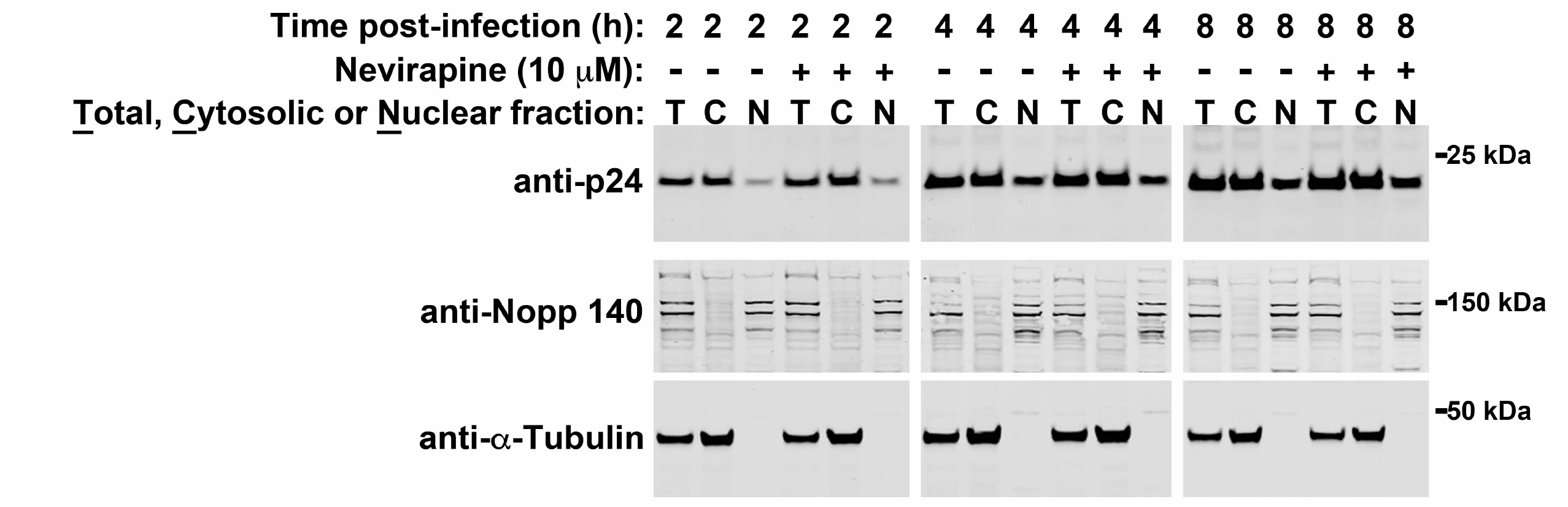
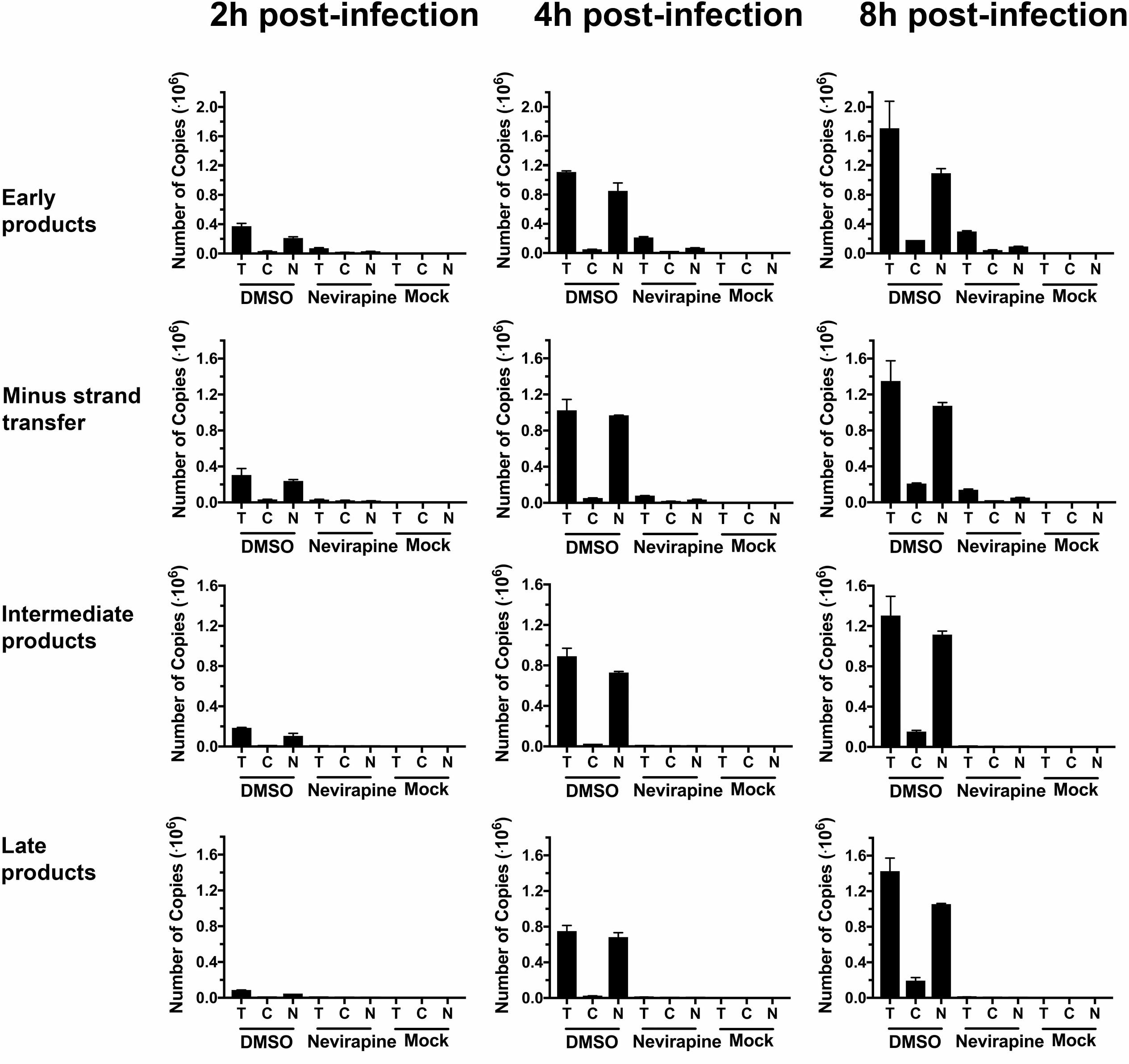
Reverse transcription occurs in the nuclear compartment. Cf2Th cells were infected with wild-type HIV-1-GFP at MOI = 2 in the presence of 10 μM nevirapine or DMSO as a vehicle control. After incubation for the indicated times, cells were fractionated and 10% aliquots of total, cytosolic, and nuclear fractions were analyzed by western blotting using anti-p24, anti-Nopp 140, and anti-tubulin antibodies **(A)** or used for DNA extraction and analyzed for the presence of HIV-1 reverse transcription intermediates (early products, minus strand transfer, intermediate products, and late products) by RT-PCR as described in methods **(B)**. Experiments were repeated three times (Figure S7) and a representative figure is shown.

## DISCUSSION

Our development of a biochemical assay to measure the amount of nuclear HIV-1 capsid during infection is based on separating HIV-1-infected cells into nuclear and cytosolic fractions for western blotting. Interestingly, we found that the amount of nuclear capsid peaks at 8 h post-infection in synchronized infections, which coincides with the peak of late reverse transcription. As a control, we used the small molecules PF74 and BI-2 that prevent 2-LTR circle formation measured by real-time PCR (Balasubramaniam et al., 2019; Fricke et al., 2014a). The absence of 2-LTR circles during HIV-1 infection is usually interpreted as a block to nuclear import, but the block may be at any step before 2-LTR circle formation, which occurs in the nucleus (Butler et al., 2001). Accordingly, we discovered that PF74 and BI-2 prevented capsid entry into the nucleus. However, it is difficult to mechanistically explain how PF74 and BI-2 prevent the nuclear import of capsid since these drugs exhibit several phenotypes, including the prevention of Nup153 and CPSF6 binding to capsid (Buffone et al., 2018; Lee et al., 2010) and accelerate HIV-1 uncoating during infection (Fricke et al., 2014a; Shi et al., 2011). Regardless of the mechanism, PF74 and BI-2 are appropriate controls for this assay since they prevent nuclear import of HIV-1 capsid. Treatment with PF74 during infection with HIV-1 virus bearing the capsid changes N57S and N74D did not prevent capsid nuclear import, which is in agreement with evidence showing that these viruses are resistant to PF74 (Buffone et al., 2018; Saito et al., 2016). Besides a specificity control, the infectivity of these mutant viruses correlates the presence of nuclear capsid with productive infection.

Recent reports using imaging have shown that the HIV-1 capsid enters the nuclear compartment; however, the role of capsid in the nucleus is not understood. These studies have shown that the pre-integration complex in the nucleus is associated with capsid (Bejarano et al., 2019; Burdick et al., 2017; Chin et al., 2015; Fassati and Goff, 2001; Francis and Melikyan, 2018; Hulme et al., 2015; Peng et al., 2014), which interacts with CPSF6. Depletion of CPSF6, or the use of HIV-1-N74D virus that does not interact with CPSF6, results in virus with changed integration sites that does not affect productive infection in cell lines (Buffone et al., 2018; Sowd et al., 2016). This evidence suggests that nuclear capsid might be playing a role in HIV-1 integration. Our experiments herein show that CPSF6 forms nuclear speckles upon HIV-1 infection, which might be important for productive infection. Intriguingly, we demonstrate that CPSF6 speckles only form when the binding of CPSF6 to capsid is intact, and when the capsid is physically in the nucleus. A possible scenario is that the pre-integration complex contains capsid in order to recruit CPSF6, which is important for integration site selection.

The presence of capsid in the nucleus during HIV-1 infection agrees with earlier findings suggesting that capsid is the determinant for nuclear import (Yamashita and Emerman, 2004; Yamashita et al., 2007), a concept strengthened by the ability of nucleopore proteins such as Nup153 to interact with capsid (Di Nunzio et al., 2013; Matreyek et al., 2013). The fact that capsid is in the nucleus raised the question of how capsid reaches the nuclear compartment. We considered that capsid might be reaching the nucleus in two different oligomeric states: assembled and/or disassembled. To this end, we took advantage of the ability of TRIM5α_rh_ and TRIMCyp to disassemble the HIV-1 core in the cytoplasm of infected cells (Diaz-Griffero et al., 2007; Stremlau et al., 2006). Remarkably, our experiments reveal that HIV-1 capsid does not reach the nuclear compartment of cells stably expressing TRIM5α_rh_ or TRIMCyp, implying that assembled capsid is necessary to enter the nuclear compartment and raising the question of what degree of capsid assembly is required for the complex to enter the nucleus. Although this is a difficult question, we can conclude that the level of capsid assembly after the core encounters TRIM5α_rh_ or TRIMCyp is not sufficient for nuclear import. One possibility is that successful nuclear import of the HIV-1 replication complex depends upon its association with assembled capsid; this assembled capsid may not be a complete core, but might need to consist of a substantial structure. The entry of assembled capsid into the nucleus is not a novel concept since it is documented that for other viruses, completely assembled capsids are imported into the nucleus. For example, during human hepatitis B virus infection, the intact assembled capsid, which is an icosahedral structure 36 nm in diameter, crosses the nuclear pore to enter the nucleus (Fay and Pante, 2015). Similarly, baculovirus, which are rod shape structures of 60-300 nm, enters intact into the nucleus using the nuclear pore (Fay and Pante, 2015). Therefore, it is not unlikely that HIV-1 uses a similar nuclear import strategy. By contrast, we found that the HIV-1 capsid reaches the nuclear compartment of cells stably expressing MxB or NES-CPSF6, which are proteins that prevent HIV-1 uncoating or stabilize the core (De Iaco et al., 2013; Fricke et al., 2013; Fricke et al., 2014b). This led us to conclude that factors that stabilize the HIV-1 core did not prevent nuclear capsid import, strengthening the notion that nuclear capsid is the result of importing assembled HIV-1 capsid.

To further understand whether assembled capsid enters the nuclear compartment, we took advantage of the capsid changes A14C/E45C, which result in the formation of disulfide bonds between capsid monomers that stabilize the HIV-1 core. Interestingly, we found that the electrophoretic migration for most of the HIV-1 capsid found in the nucleus was sensitive to reducing agent, implying that most of the capsid found in the nucleus is assembled which strengthens the notion that assembled capsid enters the nuclear compartment. Interestingly, HIV-1-A14C/E45C viruses undergo normal reverse transcription consistent with the notion that reverse transcription occurs before or during uncoating.

Several lines of evidence suggest that HIV-1 reverse transcription occurs before or during uncoating (Arfi et al., 2009; Hulme et al., 2011; Roa et al., 2011; Stremlau et al., 2006; Yang et al., 2013), which is defined as dissociation of monomeric capsids from the core. Therefore, we decided to test whether inhibiting reverse transcription modulates capsid import into the nucleus. We observed that inhibition of reverse transcription does not change the amount of capsid reaching the nucleus when compared to wild-type virus. These results suggest that capsid import into the nucleus is independent of reverse transcription. Accordingly, we observed that the reverse transcription block imposed by SAMHD1 to HIV-1 does not prevent capsid transport into the nucleus during infection(data not shown). Because SAMHD1 is a nuclear protein, it is likely that reverse transcription is inhibited in the nucleus, suggesting that it is also occurring in the nuclear compartment.

Our studies showed that assembled capsid reaches the nucleus; therefore, it is reasonable to think that at least part of the reverse transcription process is occurring in the nuclear compartment. To this end, we measured the different steps of reverse transcription in nuclear and cytosolic fractions over time. These investigations revealed that the nuclear fraction was enriched with reverse transcription intermediates when compared to the cytosolic fraction. Although the conventional view is that reverse transcription occurs in the cytoplasm followed by the transport of the complete viral DNA into the nucleus, we show that most of the reverse transcription intermediates are found in the nuclear compartment. Conceptually, the occurrence of reverse transcription in the nucleus is more efficient since virus integration occurs in the nucleus.

Combining our results with the work of others suggests a model in which the HIV-1 core enters the cytoplasm to begin a slow uncoating process. Although the HIV-1 replication complex slowly loses capsid monomers, it has to reach the nuclear pore containing a substantial amount of the assembled capsid to gain access to the nucleus. And while reverse transcription likely begins in the cytoplasm, our results indicate that most of the reverse transcription intermediates are found in the nuclear compartment, which is functionally expected since integration occurs in the nuclear compartment. With most of the reverse transcription occurring in the nucleus, and reverse transcription occurring before or during uncoating, our findings indicate that uncoating also occurs in the nucleus. These results fundamentally change our understanding of HIV-1 infection and conclude that nuclear import precedes reverse transcription and uncoating.

## ACKNOWLDGEMENTS

We thank the NIH AIDS repository for important reagents. M.P. acknowledge support from National Institutes of Health grant T32 AI07501. A.S. and F.D.-G. are supported by an NIH NIAID grant R01 AI087390 to F.D.-G.

## AUTHOR CONTRIBUTIONS

A.S., M.P., and K.L. conducted experiments. A.S., V.K. and F.D.-G. designed experiments and analyzed data. F.D.-G. wrote the manuscript.

## Methods

### Cell lines and drugs

Human MOLT3, HT1080, A549, HeLa, and 293T cells and canine Cf2Th cells obtained from the American Type Culture Collection (ATCC) were grown at 37°C in Dulbecco’s Modified Eagle Medium (DMEM) supplemented with 10% FBS and 1% penicillin-streptomycin. PF74 was obtained from Sigma Aldrich (SML0835). Nevirapine and AZT were obtained from the NIH AIDS repository program. BI-2, PF74, nevirapine, and AZT were dissolved in dimethyl sulfoxide (DMSO) to create a stock of 10 mM.

### Subcellular fractionation to detect HIV-1 capsid in the nucleus

The specified mammalian cells (5 × 10^6^ cells) were challenged with HIV-1 viruses at a MOI = 2 for the indicated times. Cells were harvested using trypsin for 2-5 minutes at 37°C. Harvested cells were washed twice with 1× cold PBS by centrifugation at 2000 rpm for 7 min at 4°C. The supernatant and pellet correspond to cytosolic and nuclear fractions, respectively. Supernatant was removed and centrifuged once more for complete removal. Cell pellets were resuspended in 500 μL lysis buffer (10 mM Tris pH = 6.8, 1 mM DTT, 1 mM MgCl_2_, 10% sucrose, 100 mM NaCl, 0.5% NP-40, 1× protease inhibitor). A 50 μL sample from the 500 μL lysis buffer was obtained to measure the *total* amount of capsid. The remaining 450 μL were incubated for 5 min on ice. Subsequently, the sample was centrifuged at 2000 rpm for 2 min at 4°C. Next, 400 μL of cytosolic fraction was mixed with 100 μL of 5× Laemmli buffer, which represents the *cytosolic* fraction. The nuclear pellet was washed using 1 mL of lysis buffer without NP-40 (10 mM Tris, pH = 6.8, 1 mM DTT, 1 mM MgCl_2_, 10% sucrose, 100 mM NaCl, 1× protease inhibitor) by gently inverting the tube 2-3 times. The sample was then centrifuged at 2500 rpm for 2 min at 4°C, and the supernatant was discarded. The nuclear pellet was then resuspended in 400 μL of extraction buffer (10 mM Tris, pH = 6.8, 1 mM DTT, 1 mM MgCl_2_, 10% sucrose, 400 mM NaCl, 1× protease inhibitor), and incubated on ice for 10 min. 400 μL of the nuclear fraction was then mixed with 100 μL of 5× Laemmli buffer, which represents the *nuclear* fraction. Proportional amounts of *total, cytosolic,* and *nuclear* fractions were analyzed by western blot using anti-p24 and anti-GAPDH antibodies detailed below.

### Creation of stable cell lines expressing different proteins

Retroviral vectors encoding rhesus TRIM5α_rh_, owl monkey TRIMCyp (Diaz-Griffero et al., 2006; Stremlau et al., 2006), human CPSF6(1-358) (Fricke et al., 2013), or human MxB proteins were created using the pLPCX vector (Fricke et al., 2014b). The different proteins were tagged with an influenza hemagglutinin (HA) or FLAG epitope tag at the C terminus. Recombinant viruses were produced in 293T cells by cotransfecting pLPCX plasmids with pVPack-GP and pVPack-VSV-G packaging plasmids (Stratagene). The pVPack-VSV-G plasmid encodes the vesicular stomatitis virus (VSV) G envelope glycoprotein, which allows efficient entry into a wide range of vertebrate cells. The indicated mammalian cells were transduced and selected in the appropriate concentration of puromycin (1-5 μg/mL).

### Production of HIV-1-GFP viruses

Recombinant HIV-1 expressing GFP were prepared as previously described (Diaz-Griffero et al., 2008). All recombinant viruses were pseudotyped with the VSV-G glycoprotein. For infections, 3 × 10^4^ human cells seeded in 24-well plates were incubated at 37°C with virus for 24 h. Cells were washed and returned to culture for 48 h, then subjected to flow cytometric analysis with a FACScan (Becton Dickinson). HIV-1 viral stocks were titrated by serial dilution on susceptible cells to determine the infectivity of viruses.

### Real-time PCR to detect reverse transcription intermediates

Total DNA was extracted from 10% aliquots of each fraction (*total, cytosolic*, and *nuclear*, obtained during subcellular fractionation at 2 h, 4 h, and 8 h post-infection) using the QIAamp DNA micro kit (QIAGEN). As a parallel control, another 10% aliquot of the same fractions were analyzed by western blotting using anti-p24, anti-Nopp 140, and anti-tubulin antibodies. Viral DNA forms of HIV-1 were amplified using real-time PCR. Reactions were performed in 1× TaqMan Universal Probe Master Mix II, with UNG 2× (Thermo Fischer Scientific) in 20 μL volume. The PCR reaction consisted of the following steps: initial denaturation (95°C for 15 min), 40 cycles of amplification (95°C for 15 sec, 58°C for 30 sec, 72°C for 30 sec). Primer or probe sequences are as follows: early products: hRU5-F2: 5’-GCCTCAATAAAGCTTGCCTTGA-3’; hRU5-R: 5’-TGACTAAAAGGGTCTGAGGGATCT-3’; hRU5-P: 5’-(FAM)-AGAGTCACACAACAGACGGGCACACACTA-(TAMRA)-3’; minus strand transfer: FST-F1: 5’-GAGCCCTCAGATGCTGCATAT-3’, SS-R4: 5’-CCACACTGACTAAAAGGGTCTGAG-3’, P-HUS-SS1: 5’-(FAM)-TAGTGTGTGCCCGTCTGTTGTGTGAC-(TAMRA)-3’; intermediate products: GagF1: 5’-CTAGAACGATTCGCAGTTAATCCT-3’, GagR1: 5’-CTATCCTTTGATGCACACAATAGAG-3’, P-HUS-103: 5’-(FAM)-CATCAGAAGGCTGTAGACAAATACTGGGA-(TAMRA)-3’; late products: MH531: 5’-TGTGTGCCCGTCTGTTGTGT-3’; MH532: 5’-GAGTCCTGCGTCGAGAGATC-3’; LRT-P: 5’-(FAM)-CAGTGGCGCCCGAACAGGGA-(TAMRA)-3’.

### Western blot analysis

Proteins were detected by western blot using anti-p24 (1:1,000 dilution, #3637, NIH), anti-Nopp 140 (1:5,000 dilution, a generous gift from U. Thomas Meier, Albert Einstein College of Medicine), anti-tubulin (1:8,000 dilution, #PA5-29444, Invitrogen), anti-FLAG (1:1,000 dilution, Sigma), anti-hemagglutinin (HA) (1:1,000 dilution, Sigma), and anti-glyceraldehyde-3-phosphate dehydrogenase (GAPDH) (1:5,000 dilution, Invitrogen). Secondary antibodies against rabbit and mouse IgG conjugated to IRDye 680LT or IRDye 800CW were obtained from Li-Cor (1:10,000 dilution). Bands were detected by scanning blots using the Li-Cor Odyssey imaging system in the 700 nm or 800 nm channels.

### Quantification and Statistical Analysis

Statistical analyses were performed using unpaired t-test. Information on sample number, replicates, and p value can be found in corresponding figure legends. Quantification of western blot band intensity was performed using ImageJ. For all experiments, the mean and standard deviation values were calculated using GraphPad Prism 7.0c.

## SUPPLEMENTAL INFORMATION

**Figure S1.**
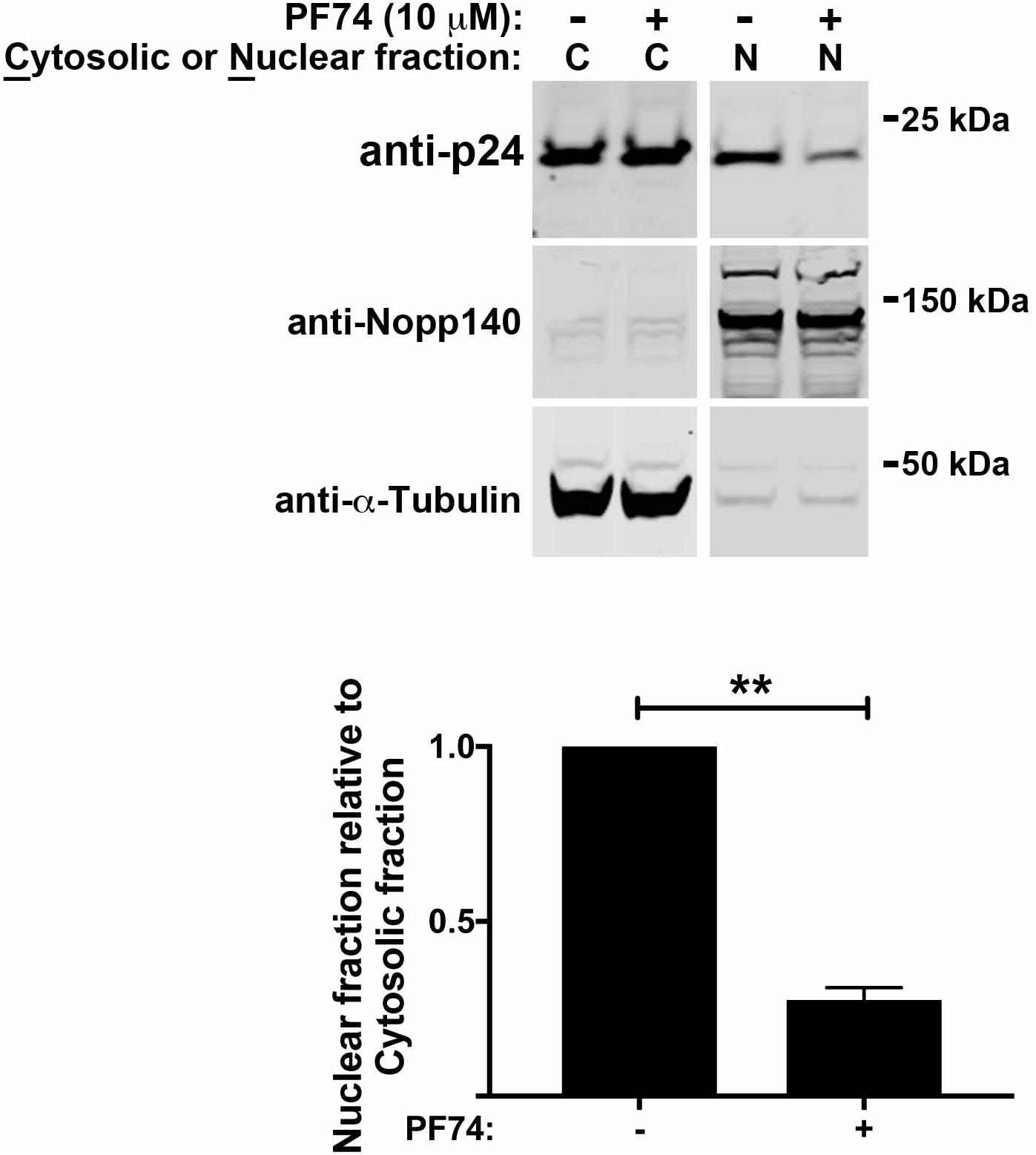
PF74 prevents the nuclear import of HIV-1 capsid in T cells. MOLT3 cells were infected with HIV-1–GFP at MOI = 2 in the presence of 10 μM PF74 or DMSO as a vehicle control for 8 h. Cells were separated into nuclear and cytosolic fractions and analyzed for capsid content by western blotting using anti-p24 antibodies. Detection of Nopp140 and tubulin by western blotting was used as a marker for nuclear and cytosolic content, respectively. Experiments were repeated at least three times and a representative figure is shown. The ratio of Nuclear to Cytosolic capsid for three independent experiments with standard deviations is shown. ** indicates P-value < 0.001 as determined by using the unpaired t-test.

**Figure S2.**
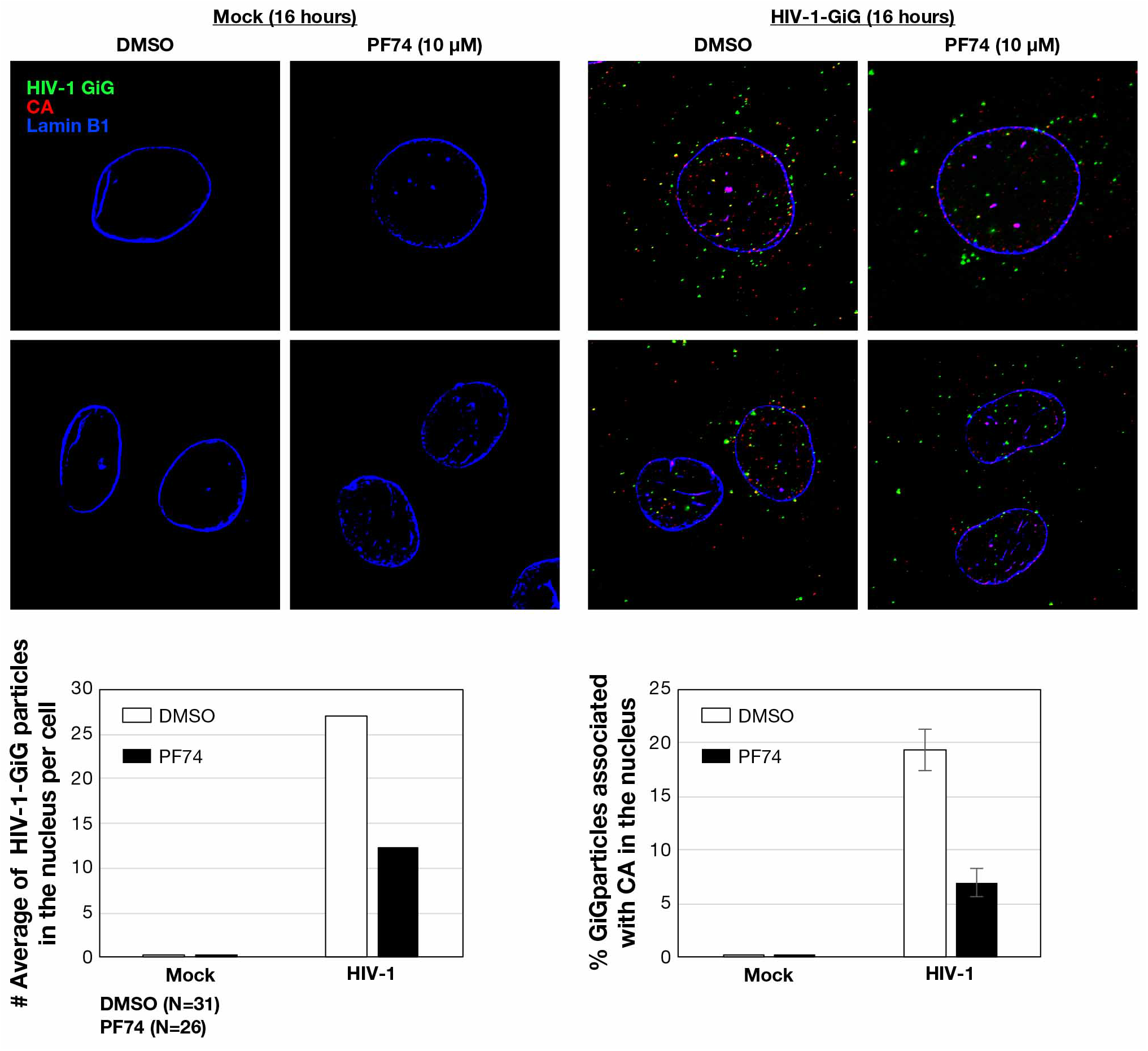
PF74 prevents the entry of capsid into the nucleus. HeLa cells were mock-infected or infected with wild-type HIV-1-Gag-protease-integrase-GFP (GiG) at MOI = ~2 in the presence of 10 μM PF74 or DMSO as a vehicle control. After incubation for 16 h, cells were fixed and stained for HIV-1-GiG, HIV-1 capsid (CA), and Lamin B1. Statistical analysis for the number of HIV-1-GiG particles in the nucleus per cell and is shown in the lower panels.

**Figure S3.**
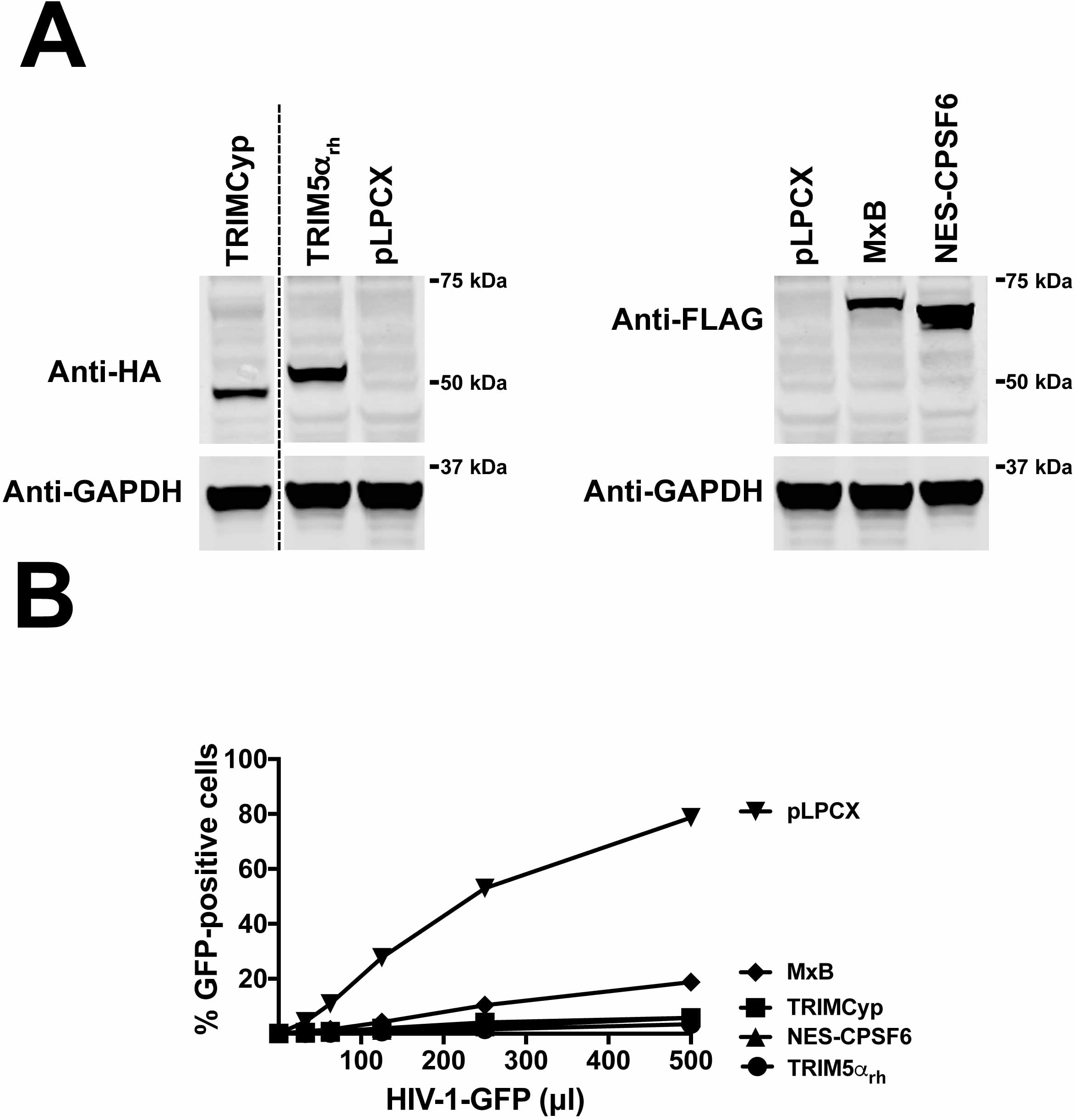
Expression of TRIM5α_rh_, TRIMCyp, CPSF6, and MxB in human cells blocks HIV-1 infection. Human HT1080 cells stably expressing the indicated restriction factors or proteins **(A)** were challenged with increasing amounts of HIV-1-GFP for 48 h. Subsequently, infectivity was determined by measuring the percentage of GFP-positive cells using a flow cytometer **(B)**. Experiments were repeated at least three times and a representative figure is shown.

**Figure S4.**
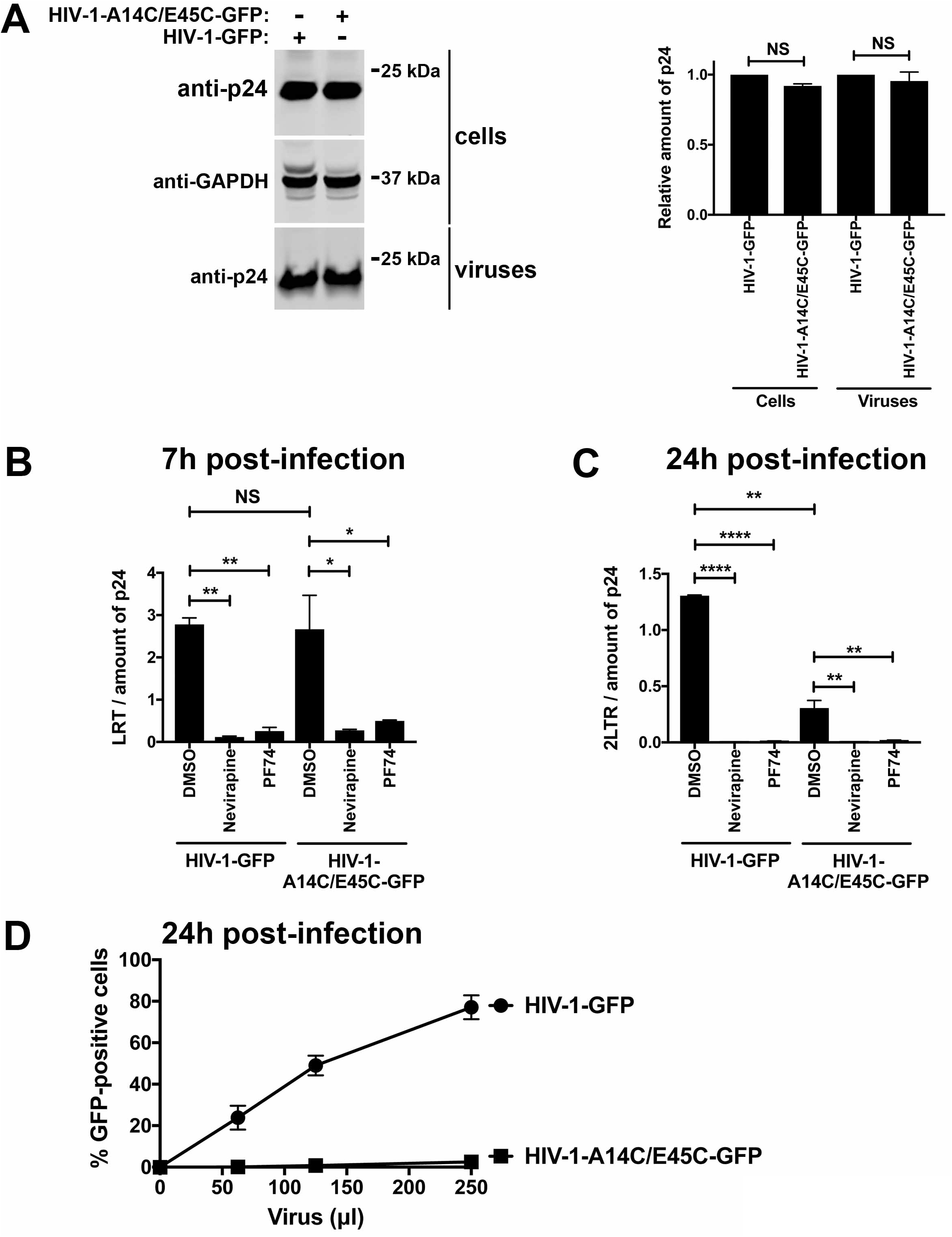
Mutant HIV-1-A14C/E45C viruses undergo normal reverse transcription, but are defective in the formation of 2-LTR circles and productive infection. **(A)** Mutant HIV-1-A14C/E45C viruses did not show a defect on particle release and maturation when compared to wild type HIV-1. Wild type and mutant HIV-1 viruses were produced by transfection of appropriate plasmids in human 293 T cells, as described in Methods. 48 hours post-transfection cells and supernatants were collected. Cell lysates and concentrated supernatants on a 20% sucrose cushion were analyzed by Western blotting using anti-p24 antibodies. **(B)** Mutant HIV-1-A14C/E45C viruses undergo normal reverse transcription when compared to wild type HIV-1. Human A549 cells were challenged with p24-normalized viruses and used to measure late reverse transcripts 7 hours post-infection by RT-PCR, as described in Methods. **(C)** Mutant HIV-1-A14C/E45C viruses are defective in the formation of 2-LTR circles when compared to wild type HIV-1. Human A549 cells were challenged with p24-normalized viruses and used to measure the formation of 2-LTR circles 24 hours post-infection by RT-PCR, as described in Methods. * indicates P-value < 0.005, ** indicates P-value < 0.001, **** indicates P-value < 0.0001, NS indicates not significant as determined by using the unpaired t-test. **(D)** Mutant HIV-1-A14C/E45C viruses showed a defect in achieving productive infection. Increasing amounts of wild type and mutant HIV-1 viruses were used to challenge A549 cells for 48 h. Subsequently, infectivity was determined by measuring the percentage of GFP-positive cells using a flow cytometer.

**Figure S5.**
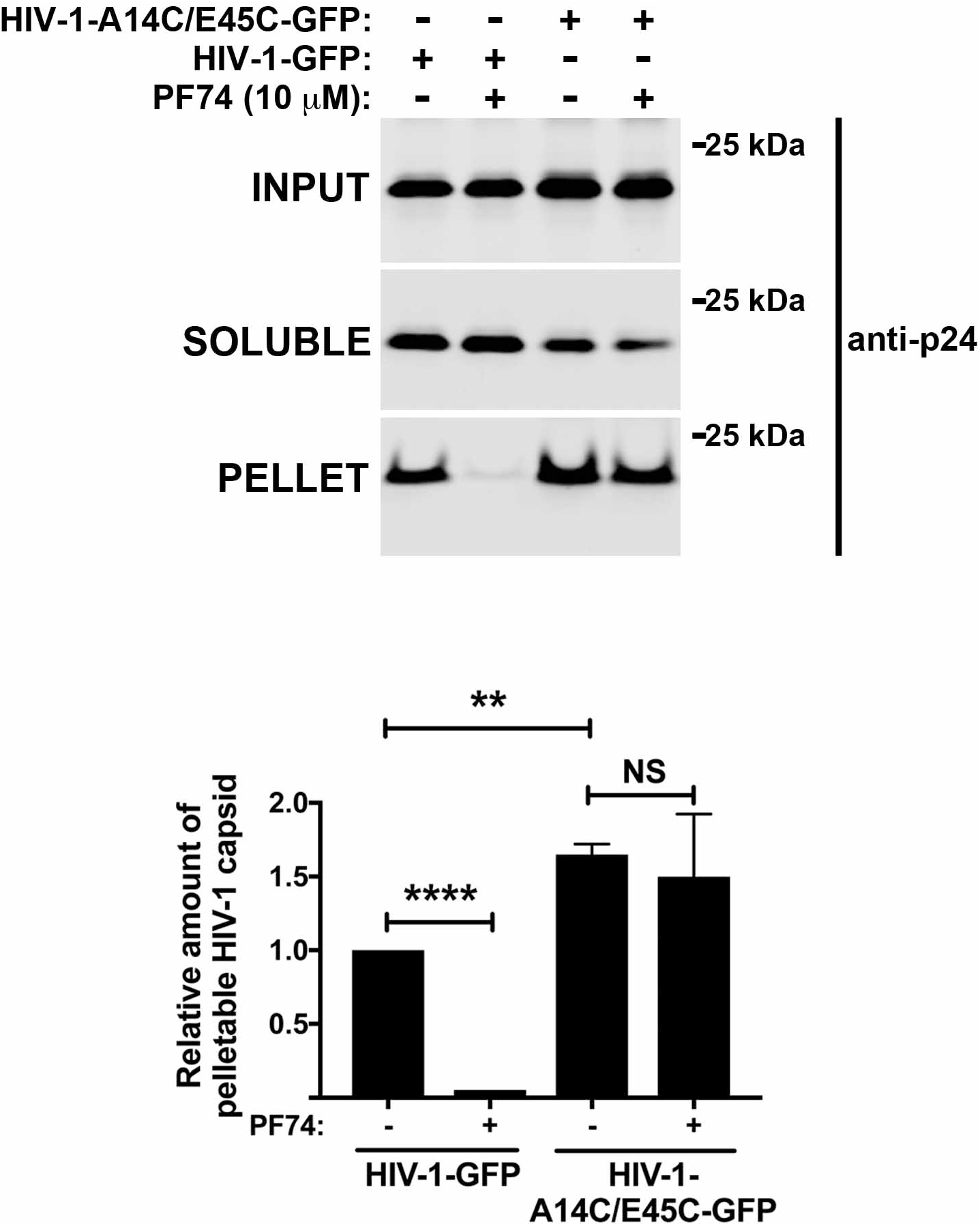
HIV-1 viruses bearing capsid changes A14C/E45C showed greater capsid stability when compared to wild-type. Human A549 cells were incubated with p24-normalized amounts of HIV-1-GFP and HIV-1-A14C/E45C viruses in the presence of 10 μM PF74 at 4°C for 30 min. Cells were washed and returned to 37°C, and infection was allowed to proceed for 12 h in the presence of 10 μM PF74 or DMSO as vehicle control. Cell extracts were fractionated on sucrose gradients as described in Methods. Input, soluble, and pellet fractions were analyzed by western blotting using antibodies against the HIV-1 p24 capsid protein. Experiments were repeated at least three times and a representative figure is shown. The fraction of pelletable capsid for three independent experiments with standard deviations is shown. ** indicates P-value < 0.001, **** indicates P-value < 0.0001, NS indicates not significant as determined by using the unpaired t-test.

**Figure S6.**
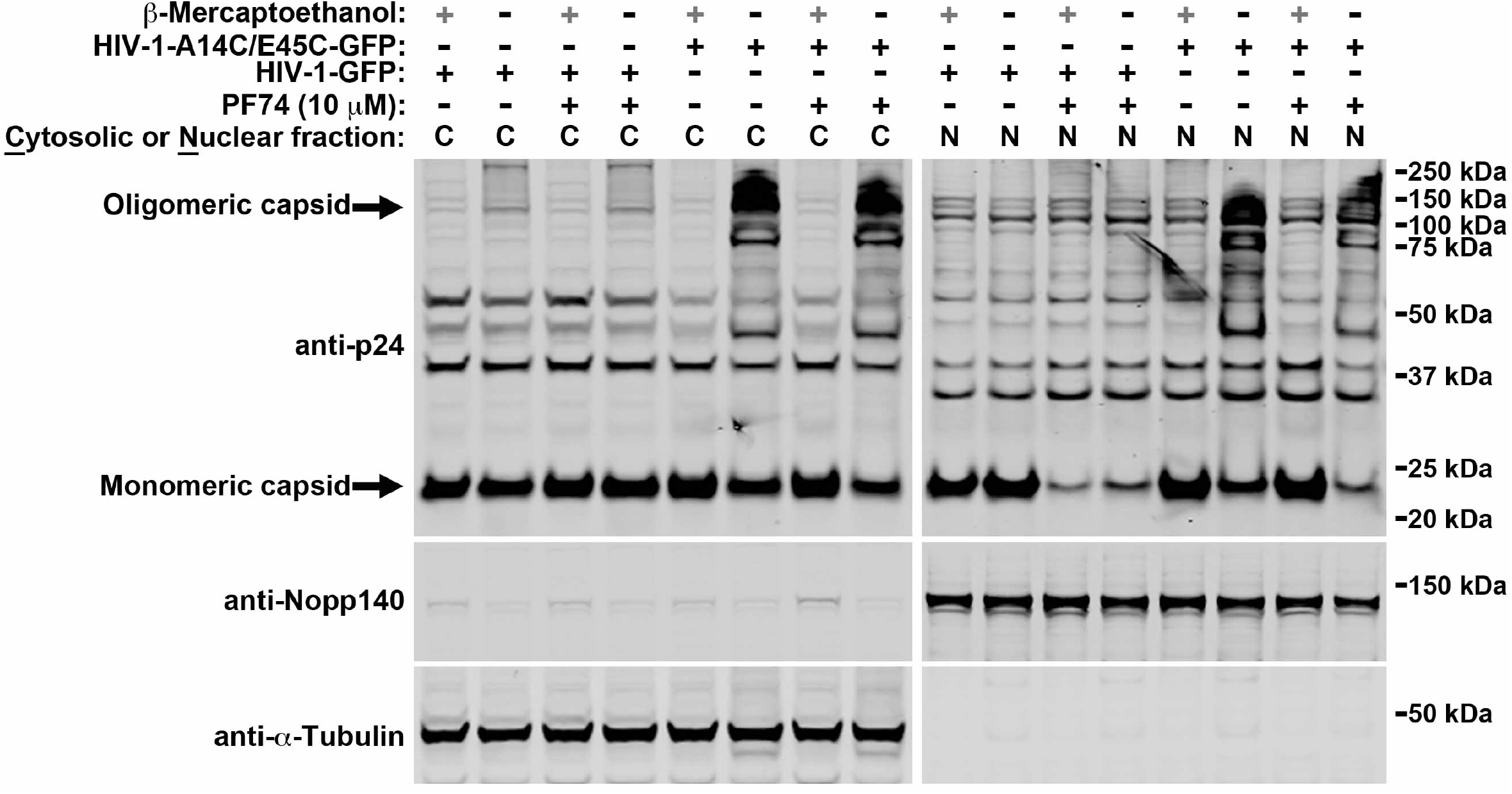
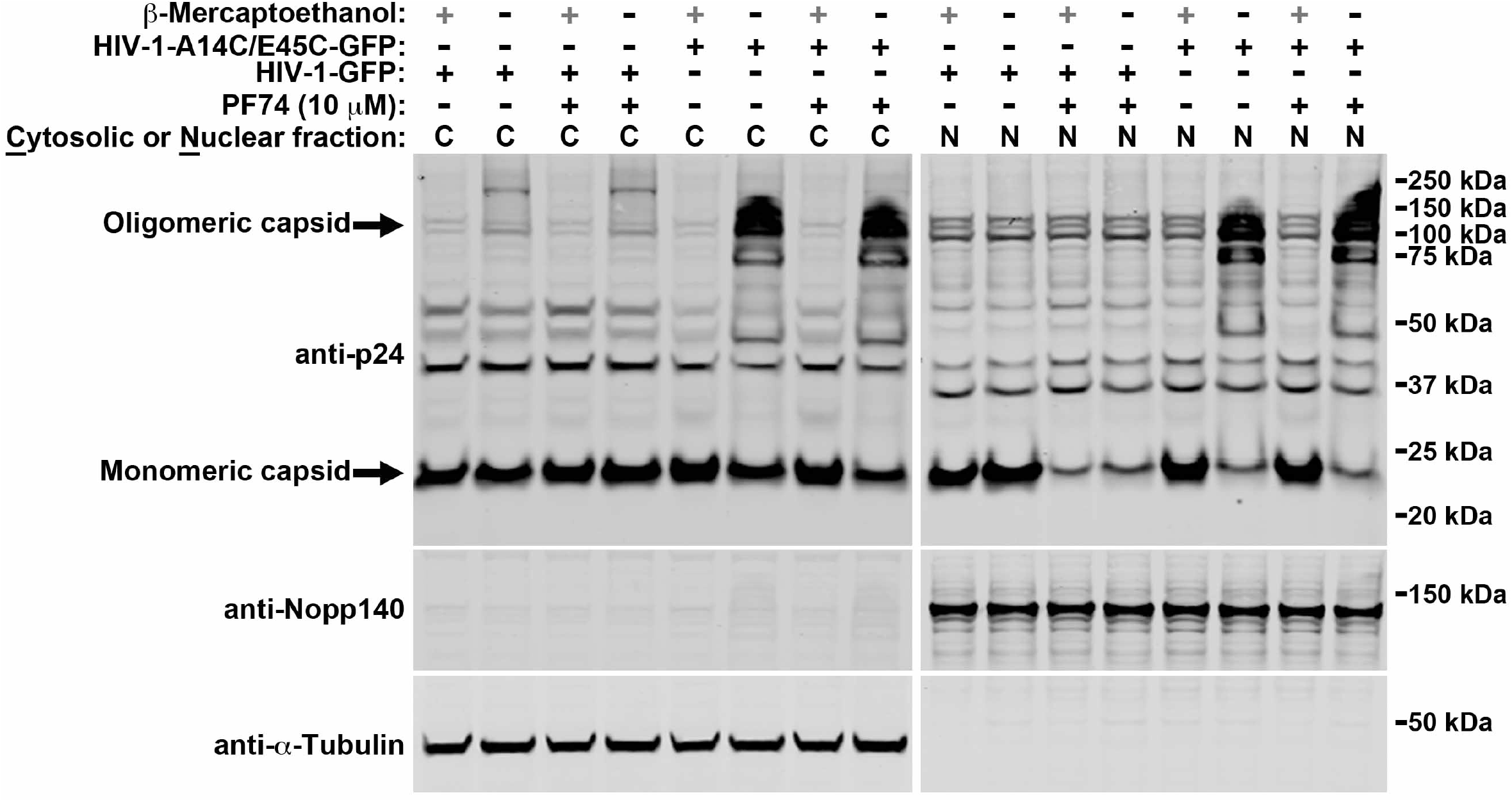
Assembled capsid complexes are imported into the nuclear compartment during HIV-1 infection. **(A, B)** Human A549 cells were infected using p24-normalized amounts of wild-type HIV-1-GFP and mutant HIV-1-A14C/E45C-GFP viruses (virus amount corresponded to wild-type MOI = 2) in the presence of 10 μM PF74 or DMSO as a vehicle control for 8 h. Cells were separated into nuclear and cytosolic fractions and analyzed for capsid content by western blotting using anti-p24 antibodies in the presence or absence of the reducing agent β-mercaptoethanol. Detection of Nopp140 and tubulin by western blotting was used as a marker for nuclear and cytosolic content, respectively.

**Figure S7.**
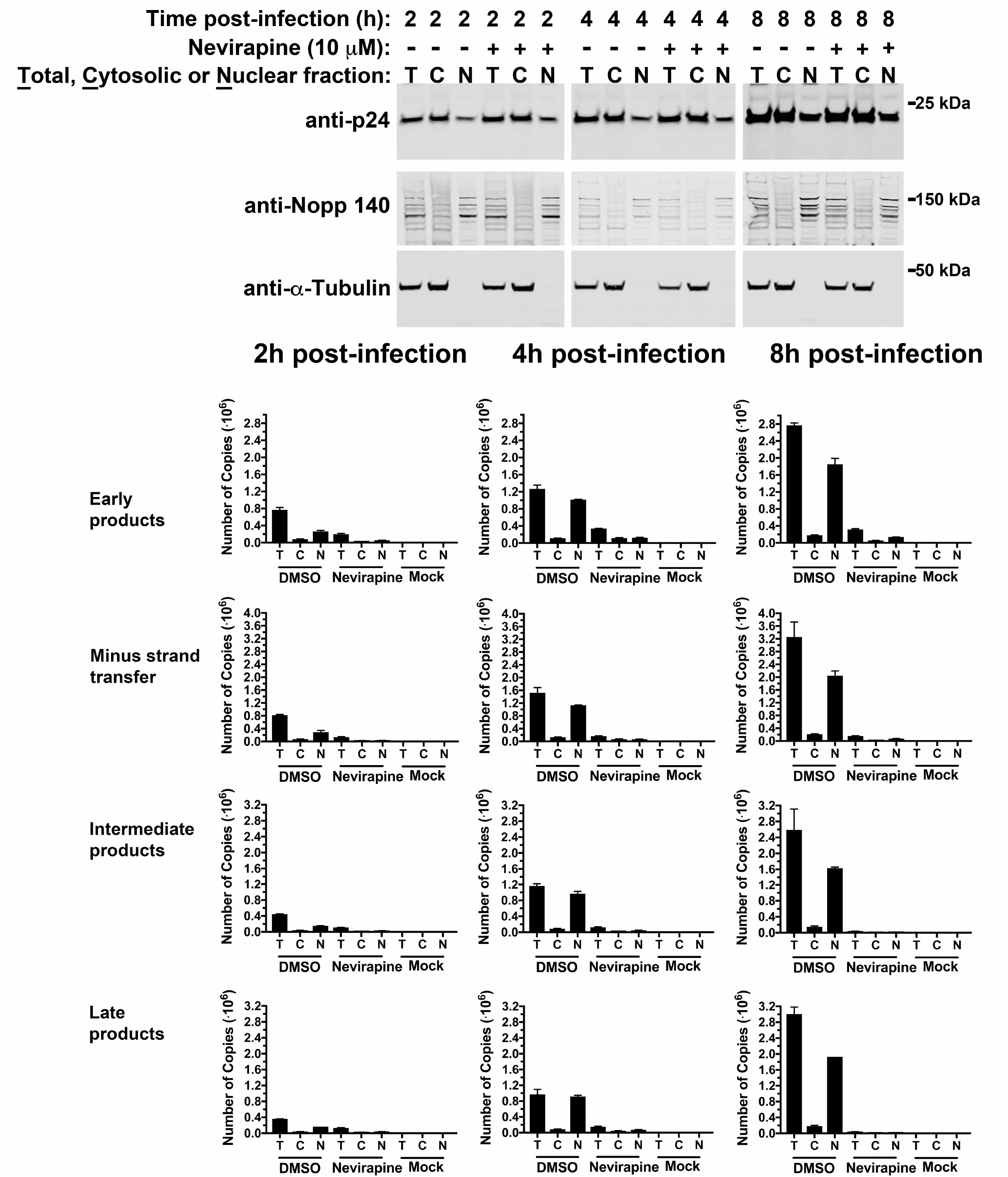
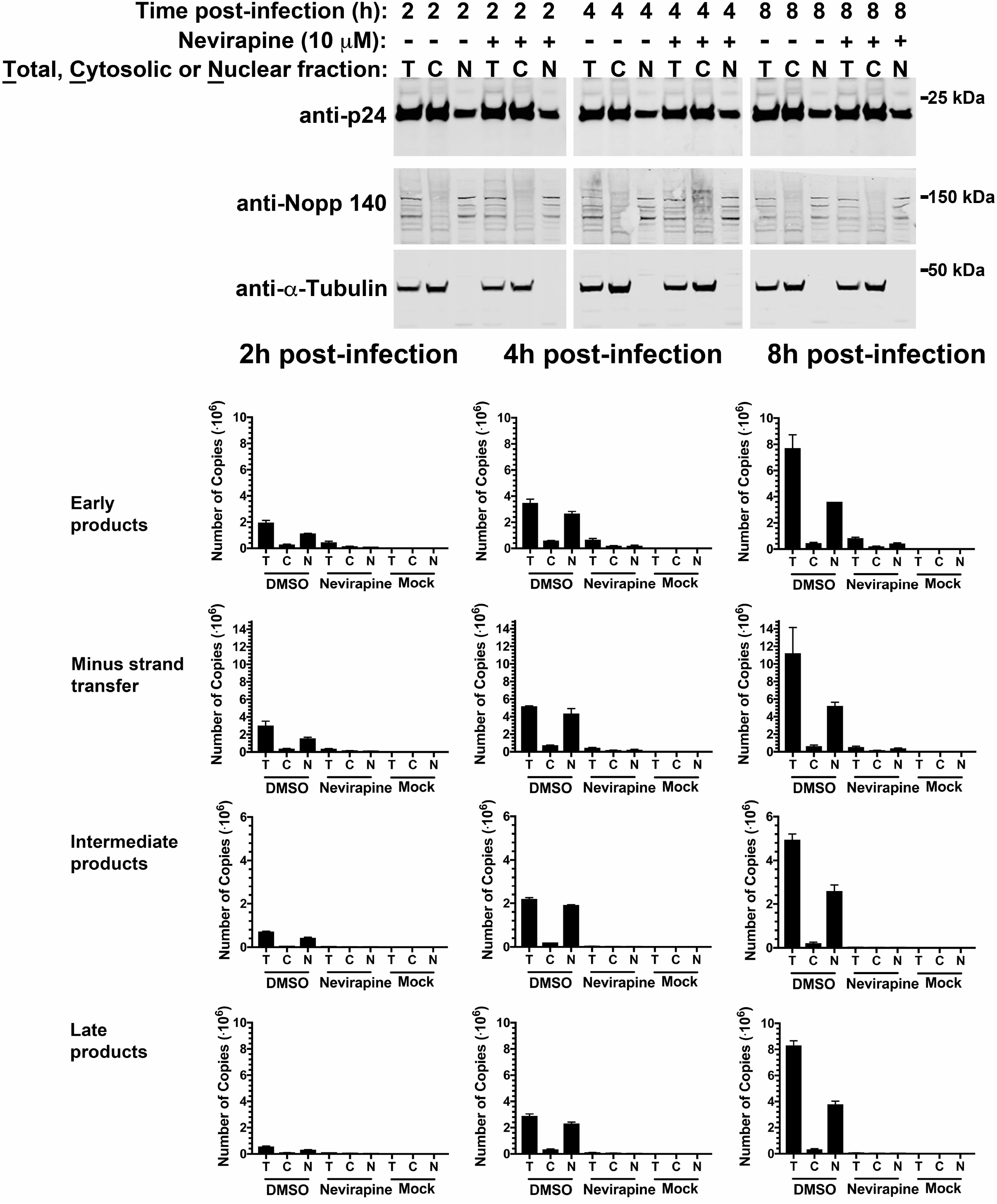
Reverse transcription occurs in the nuclear compartment. **(A, B)** Human cells were infected with wild-type HIV-1-GFP at MOI = 2 in the presence of 10 μM nevirapine or DMSO as a vehicle control. After incubation for the indicated times, cells were fractionated and 10% aliquots of total, cytosolic, and nuclear fractions were analyzed by western blotting using anti-p24, anti-Nopp 140, and anti-tubulin antibodies or used for DNA extraction and analyzed for the presence of HIV-1 reverse transcription intermediates (early products, minus strand transfer, intermediate products, and late products) by RT-PCR as described in methods.

